# Sequential defects in cardiac lineage commitment and maturation cause hypoplastic left heart syndrome

**DOI:** 10.1101/2021.04.24.441110

**Authors:** Markus Krane, Martina Dreßen, Gianluca Santamaria, Ilaria My, Christine M. Schneider, Tatjana Dorn, Svenja Laue, Elisa Mastantuono, Riccardo Berutti, Hilansi Rawat, Ralf Gilsbach, Pedro Schneider, Harald Lahm, Sascha Schwarz, Stefanie A. Doppler, Sharon Paige, Nazan Puluca, Sophia Doll, Irina Neb, Thomas Brade, Zhong Zhang, Claudia Abou-Ajram, Bernd Northoff, Lesca M. Holdt, Stefanie Sudhop, Makoto Sahara, Alexander Goedel, Andreas Dendorfer, Fleur V.Y. Tjong, Maria E. Rijlaarsdam, Julie Cleuziou, Nora Lang, Christian Kupatt, Connie Bezzina, Rüdiger Lange, Neil E. Bowles, Matthias Mann, Bruce Gelb, Lia Crotti, Lutz Hein, Thomas Meitinger, Sean Wu, Daniel Sinnecker, Peter J. Gruber, Karl-Ludwig Laugwitz, Alessandra Moretti

## Abstract

**Background:** Complex molecular programs in specific cell lineages govern human heart development. Hypoplastic left heart syndrome (HLHS) is the most common and severe manifestation within the spectrum of left ventricular outflow tract obstruction defects occurring in association with ventricular hypoplasia. The pathogenesis of HLHS is unknown, but hemodynamic disturbances are assumed to play a prominent role.

**Methods:** To identify perturbations in gene programs controlling ventricular muscle lineage development in HLHS, we performed: i) whole-exome sequencing of 87 HLHS parent-offspring trios, ii) nuclear transcriptomics of cardiomyocytes from ventricles of 4 patients with HLHS and 15 controls at different stages of heart development, iii) single cell RNA sequencing and iv) 3D modeling in iPSCs from 3 patients with HLHS and 3 controls.

**Results:** Gene set enrichment and protein network analyses of damaging *de-novo* mutations and dysregulated genes from ventricles of patients with HLHS suggested alterations in specific gene programs and cellular processes critical during fetal ventricular cardiogenesis, including cell-cycle and cardiomyocyte maturation. Single-cell and 3D modeling with iPSCs demonstrated intrinsic defects in the cell-cycle/UPR/autophagy hub resulting in disrupted differentiation of early cardiac progenitor lineages leading to defective cardiomyocyte-subtype differentiation/maturation in HLHS. Additionally, premature cell-cycle exit of ventricular cardiomyocytes from HLHS patients prevented normal tissue responses to developmental signals for growth leading to multinucleation/polyploidy, accumulation of DNA damage, and exacerbated apoptosis, all potential drivers of left ventricular hypoplasia in absence of hemodynamic cues.

**Conclusions:** Our results highlight that despite genetic heterogeneity in HLHS, many mutations converge on sequential cellular processes primarily driving cardiac myogenesis, suggesting novel therapeutic approaches.

## INTRODUCTION

HLHS is a severe form of congenital heart disease (CHD) characterized by underdevelopment of left-sided cardiac structures, including left ventricular (LV) hypoplasia, hypoplastic ascending aorta, intact interventricular septum, aortic and/or mitral valve atresia/stenosis in the setting of concordant ventriculoarterial connections^1–3^. Although a genetic etiology is supported by an increased recurrence risk and familial clustering, the largely sporadic occurrence suggests a complex genetic model^4, 5^.

Historically, ventricular and aortic hypoplasia in HLHS has been attributed to reduced growth arising as consequence of restricted blood flow due to maldeveloped mitral and/or aortic valve^6, 7^. Recently, an anatomic study of HLHS hearts indicated that ventricular and valvular morphology are poorly correlated and three phenotypic LV variants (“slit-like”, “thickened”, and “miniature”) are present, extending earlier suggestions that a pathogenic mechanism based upon reduced blood flow alone may be insufficient^8, 9^. In the *Ohia* mouse model specific mutations drive LV hypoplasia by perturbing cardiomyocyte (CM) proliferation/differentiation^10^. The large majority of *Ohia* mutants display HLHS, although some mutants can have double-outlet right ventricle that do not uniformly resemble human HLHS pathology. Interestingly,, transcriptional alterations found in the LV of the *Ohia* mice with HLHS are present, though less severe, in the right ventricle (RV). This finding suggests that intrinsic defects in myogenic programs of both ventricles may manifest differently, depending on combinations of the cardiac progenitor (CP) population affected and physiological milieu.

During cardiogenesis, two main CP lineages provide CMs to the developing heart with distinct temporal and spatial contributions^11^. The first heart field (FHF) CPs are fated to differentiate early forming the primitive heart tube, the LV, and portion of the atria, while the second heart field (SHF) cells show delayed differentiation into CMs and represent initially a reservoir of multipotent CPs^12, 13^. Later, SHF CPs give rise to the RV, the proximal outflow tract (OFT), and part of the atria^14^.

Elegant work using CMs derived from human iPSCs has recently begun to elucidate molecular mechanisms of CHD in patient-specific *in-vitro* models of cardiogenesis^15, 16^. Studies in iPSC-CMs from HLHS patients carrying variants in the NOTCH signaling pathway demonstrated impaired differentiation^17–19^; other studies implicated transcriptional repression of *NKX2-5*, *HAND1*, and *NOTCH1* or activation of atrial gene programs^20, 21^. Still lacking is a unifying picture of how the profound genetic heterogeneity of HLHS converges in common perturbations of sequential cellular processes driving heart morphogenesis, how these processes are altered, and how such alterations contribute to the disease.

Here, we combined whole-exome sequencing (WES) of parent-offspring trios, transcriptome profiling of CMs from ventricular biopsies, iPSC-derived CP/CM models of 2D/3D cardiogenesis, and single-cell gene expression analysis to decode the cellular and molecular principles of HLHS phenotypes. Our results show that initial aberrations in the cell-cycle/UPR/autophagy hub lead to disrupted CP lineage commitment. Consequently, impaired maturation of ventricular CMs (vCMs) limits their ability to respond to growth cues resulting in premature cell-cycle exit and increased apoptosis under biomechanical stress in 3D heart structures. Together, these studies provide evidence that HLHS pathogenesis is not exclusively of hemodynamic origin, and reveal novel potential nodes for rational design of therapeutic interventions.

## METHODS

### Data availability

All data needed to evaluate the conclusions of this study are present in the article or the Data Supplement. The raw omics data have been deposited at public available databases (bulk and scRNAseq, #GSE135411; nRNAseq, #PRJNA353755; proteomics, #PXD014812).

### Patients and controls

All HLHS patients harbored hypoplastic LV and ascending aorta, mitral and/or aortic valve atresia/stenosis, and intact interventricular septum. Two additional patients with LV hypoplasia and a ventricular septal defect not meeting strict HLHS criteria were included in the WES analysis but not selected for further analyses.

Blood for WES was collected from 87 probands with sporadic HLHS and their respective parents (clinical information are summarized in Table I Supplement). Human heart ventricular samples from control and HLHS patients were obtained from aborted fetuses or during the Norwood Stage-I palliation. All donors or their legal representatives provided informed consent. The study was performed according to the Declaration of Helsinki and approved by the local ethics committees at the respective institutions (KaBi-DHM: 5943/13, 247/16S; IHG: 5360/13; HSM:15-00696).

### Statistics

Statistical analyses were performed using R and Graphpad Prism. Data were analyzed with ANOVA, Kruskal-Wallis, Mann–Whitney, Chi-square, Fisher’s, and Student’s t test, as appropriate. P values and p adjusted of less than 0.05 were considered statistically significant.

## RESULTS

### Damaging *de-novo* mutations in HLHS patients are linked to cardiac development, chromatin organization, and cell-cycle phases

WES from 87 patients with sporadic HLHS and their parents recovered 94 non-synonymous (NS) *de-novo* mutations (DNMs), of which 12 were loss-of-function (LOF) and 76 predicted damaging missense (D-Mis) variants (Table I Supplement). One to five NS-DNMs were found in 56 cases (Figure IA and IB Supplement). Among the 51 cases with D-DNMs, 34 carried mutations in genes or HGNC gene families that harbored multiple hits (Figure 1A). D-DNMs in *MYRF*, *MACF1*, and *LRP6* were found twice, with 2 individuals carrying variants in 2 of the 3 genes (Figure 1A). *MYRF* was recently associated with CHD^22^ and HLHS^23^. *LRP6* and *MACF1* are linked to WNT signaling^24, 25^. Gene-families with multihits were likewise related to pivotal signaling pathways implicated in cardiac development (Hedgehog, FGF, and NOTCH) or to the histone modification H3K4me-H3K27me pathway that has been associated with CHD^26, 27^.

**Figure 1.**
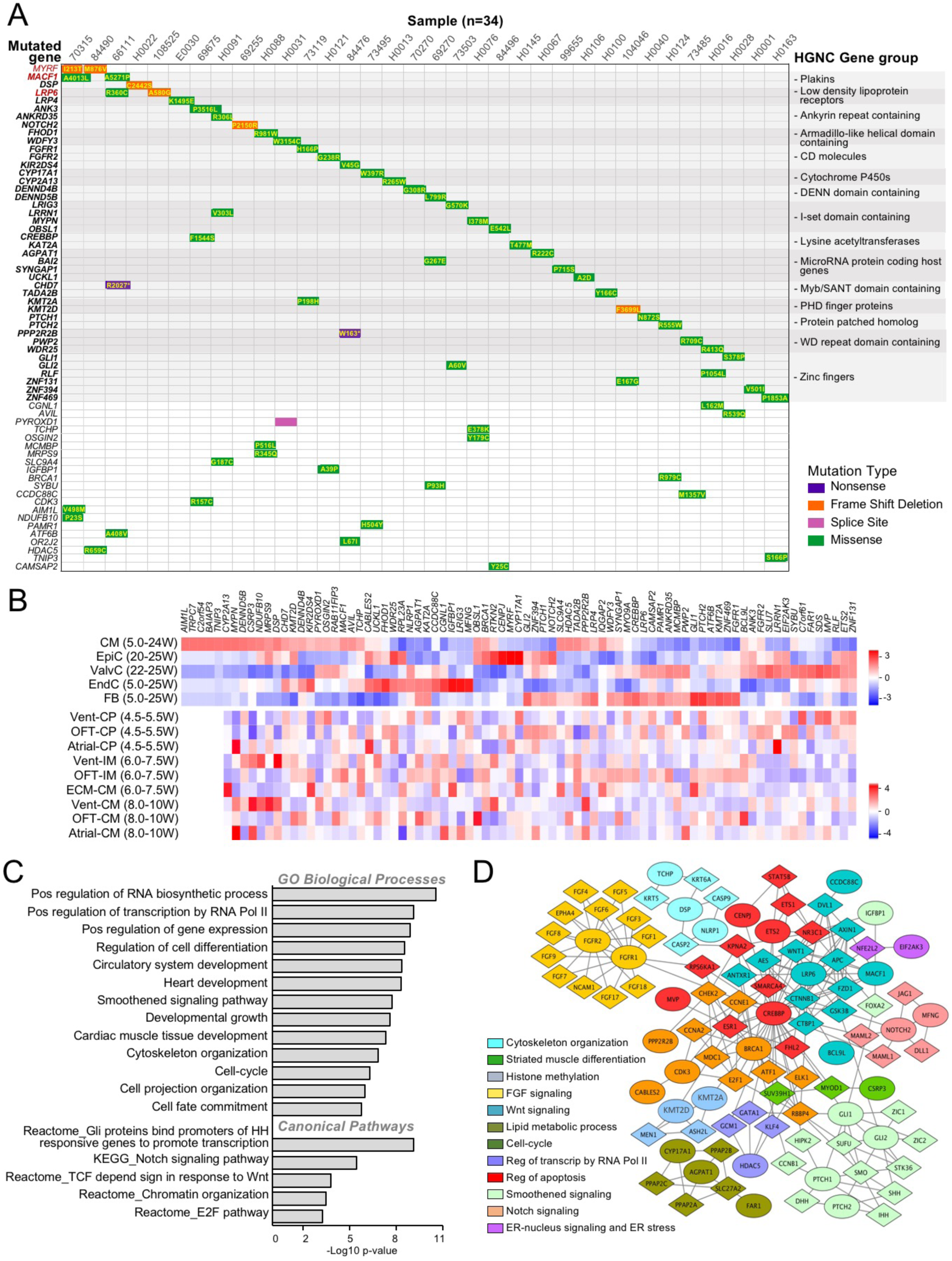
Characterization of D-DNMs in HLHS patients. **A**, Waterfall plot of de-novo multihit genes (red) or gene family (bold) identified in the HLHS cohort. Additional de-novo genes in each subject are in plain black. **B**, Cell type-specific expression of HLHS D-DNM genes based on scRNA-seq from normal fetal hearts between 4.5 and 25 weeks (W) of gestation. Data from Ciu et al.^29^ and Sahara et al.^28^ were used to generate the upper and lower heatmap, respectively. ECM, extracellular matrix; EndC, endothelial cells; EpiC, epicardial cells; FB, fibroblasts; IM, intermediates; ValvC; valvular cells; Vent, ventricular. **C** and **D**, Bar chart of Gene Ontology (GO) enrichment analysis (C) and protein-protein network analysis (D) of D-DNM genes. In D, each Netbox module is coded by a different color, with mutated genes illustrated as circle and linker genes as diamonds.

Analyzing single-cell transcriptomics from human embryonic (4.5–25 weeks post-conception) cardiac cells^28, 29^, we found 96% of the 84 D-DNM genes expressed in different cell types, significantly more than random gene sets (76.7%±8.8%, *p*-value <10e-15). Interestingly, high expression was observed in at least one stage of chamber or OFT development (Figure 1B and Figure I Supplement), suggesting an intrinsic CM lineage dysfunction as a possible determinant of HLHS. GSEA of D-DNM genes revealed a significant enrichment in gene categories related to cardiac and embryonic development/growth, cell fate commitment/differentiation as well as cell-cycle and G1/S phase transition (E2F pathway) (Figure 1C and Table I Supplement). Network analysis identified 12 interconnected protein-protein interaction modules (0.63 modularity; 11.84 scaled-modularity) significantly different from random networks (0.36±0.023 modularity (n=100, mean±SD); *p*-value <10e-05), comprising 32 of the 84 D-DNM genes (Figure 1D). Genes in these modules act in biological processes critical for CP specification and CM maturation, including cell-cycle and apoptosis, response to endoplasmic reticulum (ER) stress, signaling by Smoothened, WNT, FGF, and NOTCH, transcriptional regulation, and histone methylation.

Additionally, we interrogated independent 459 HLHS parents-offspring trios within the CHD cohort of the Pediatric Cardiac Genomics Consortium (PCGC)^27^ for *de-novo* variants. The occurrence of NS-DNMs and the distribution of variant classes were comparable, with 3 genes related to known syndromic forms of CHD (*CHD7*, *KMT2D*, and *MYRF*) being common D-DNM multihits (Figure IIA through IIC Supplement). GSEA and network interactome analyses of the combined cohort yielded similar results to our cohort alone (Figure IID and IIE Supplement). Importantly, compared to 1789 control trios comprising parents and unaffected siblings of autism probands^30^, most GSEA retrieved pathways showed significant burden enrichment in D-DNMs among cases (Figure IID Supplement). Randomization of DNMs between HLHS and controls confirmed the non-stochastic nature of our results (Table I Supplement).

### Dysregulated genes in infant HLHS CMs belong to dynamic transcriptional networks during cardiogenesis

To verify dynamic expression of HLHS D-DNM genes during CM development and identify temporal/spatial processes of cardiogenesis critical to HLHS pathogenesis, we performed nuclear RNA sequencing (RNAseq) of RV CMs from healthy hearts at 3 developmental stages – *fetal* (16-23 weeks post-conception), *infancy* (10 weeks to 10 months), and *adulthood* (21-50 years) – and from infant HLHS subjects (5-11 days of age) (Figure 2A and Table II Supplement). The choice of RV tissue was based on the fact that LV is inaccessible from living HLHS subjects and that both ventricles may share altered transcriptional signatures, as found in *Ohia* mutants.

**Figure 2.**
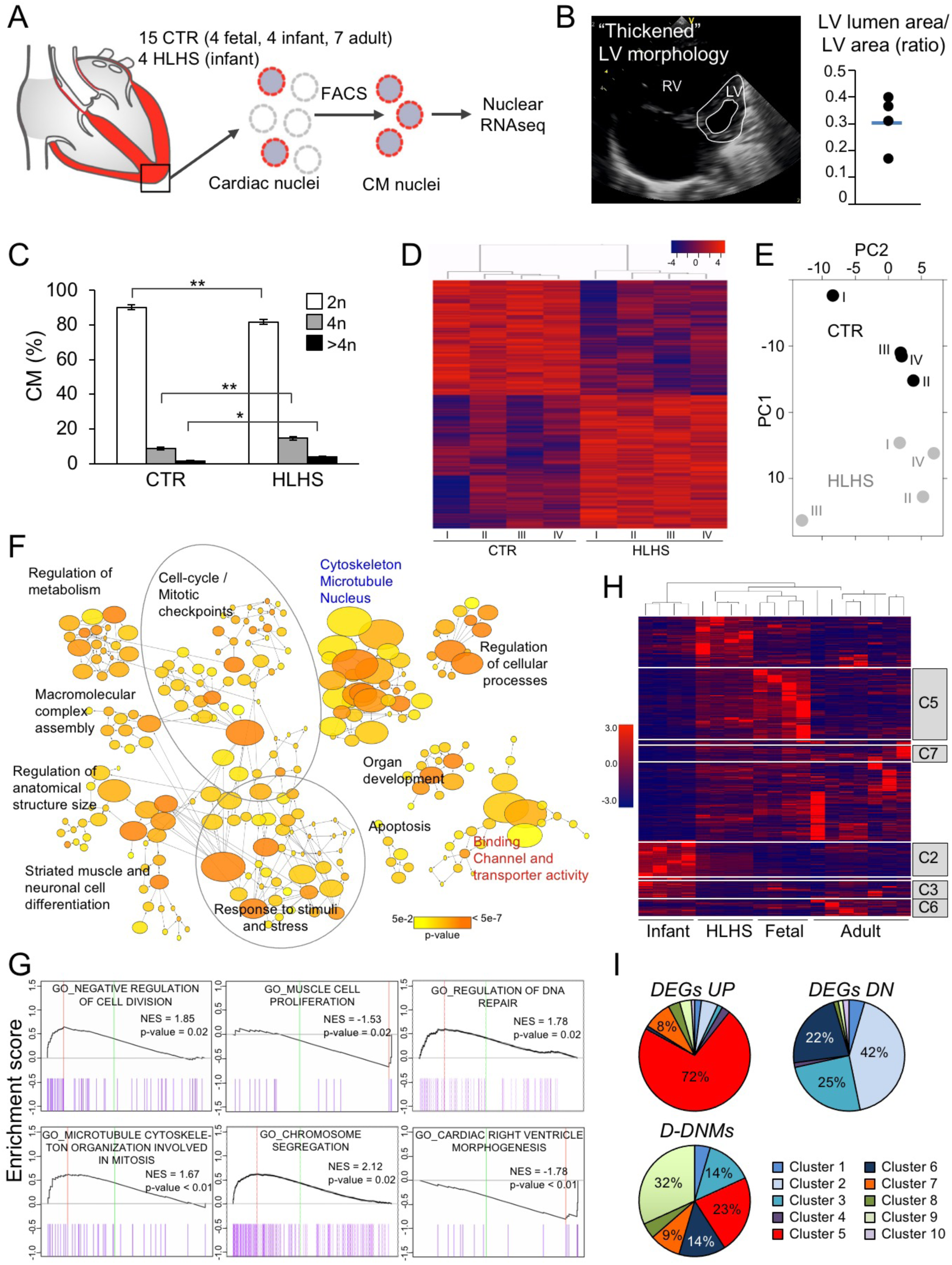
Gene expression analysis of CM nuclei from HLHS and control hearts. **A**, Workflow of CM nuclei isolation for RNAseq. **B**, Representative echocardiogram of HLHS patients with a distinct LV phenotype. Dot plot shows the ratio of LV lumen area/LV area for the 4 HLHS patients and the average value (blue line). **C**, Ploidy level of CM nuclei in HLHS and control (CTR) hearts. Data are mean ± SEM. *p<0.05, **p<0.01 (t test). **D**, Heatmap depicting normalized RNAseq expression values of DEGs (1.5-fold-expression, *p*-value ≤0.05). Gene regulations are reported as a color code and hierarchical clustering result as a dendrogram. **E**, PCA performed on rlog-normalized (DESeq2) counts for all nuclear RNAseq samples. **F**, Network visualization of the enriched GOs of HLHS DEGs using the Cytoscape plugins BinGO. Nodes represent enriched GO terms, node size corresponds to the gene number and color intensity to the *p*-value. Edges represent GO relation of Biological Process (black), Molecular Function (red), and Cellular Component (blue). **G**, Representative GSEA enrichment plots. Normalized enrichment score (NES) and *p*-value are specified. **H**, Heatmap illustrating the expression of HLHS DEGs during fetal, infant, and adult stages of normal cardiac development. The dendrogram shows clustering of the HLHS infant samples with control fetal samples. Genes belonging to developmentally regulated gene clusters from Figure IIIC in the Data Supplement are highlighted. **I**, Pie charts showing the percentage of HLHS upregulated (DEGs UP), downregulated (DEGs DN), and damaging *de-novo* affected (D-DNM) genes belonging to the developmentally regulated gene clusters from Figure IIIC in the Data Supplement.

We first compared RV tissues from the 4 normal and 4 HLHS infants, who presented “thickened” morphological features of the hypoplastic LV^8, 9^ (Figure 2B). An equal proportion of CM nuclei was found in both groups (Figure IIA Supplement). HLHS CMs showed a significant increase in the number of tetraploid nuclei at the expense of diploid nuclei (Figure 2C). Since polyploidization accompanies terminal CM differentiation and permanent cell-cycle withdrawal^31^, this suggests premature cell-cycle arrest in HLHS. RNAseq analysis revealed 2,286 differentially expressed genes (DEGs) in HLHS compared to controls (Figure 2D and Table II Supplement). Comparative principal component analysis (PCA) clearly separated HLHS and controls (Figure 2E). Enrichment Map of the Gene Ontology (GO) categories recovered from DEGs demonstrated high interconnection of functionally related gene sets associated with cell-cycle/mitotic-checkpoints, response to stimuli/stress, organ development, cell differentiation/apoptosis, cytoskeleton/microtubule/nucleus, and regulation of metabolism (Figure 2F and Table II Supplement). Importantly, most pathways overlapped with the modules recovered from the HLHS D-DNM genes. In HLHS, GSEA revealed reduced expression of genes important for cardiac ventricle morphogenesis and increased expression of genes controlling DNA repair (Figure 2G). Moreover, negative regulators of cell division were upregulated, while genes associated with muscle cell proliferation were down, supporting the notion of premature cell-cycle exit.

To assess whether DEGs and D-DNM genes in HLHS are dynamically regulated during development, we next established a global expression atlas of *fetal*, *infant* and *adult* healthy RV genes. Hierarchical clustering demonstrated distinct expression profiles among developmental stages (Figure IIIB Supplement). Investigation of DEGs identified 10 transcriptome clusters dynamically changing during cardiac development (Figure IIIC Supplement). DEGs in HLHS infants resembled the *fetal* rather than the *infant* stage (Figure 2H). Remarkably, 60% of HLHS downregulated DEGs belonged to dynamic transcriptome clusters, being 56% distributed in gene clusters 1, 2, 3, and 6. These contained genes upregulated from the embryonic to newborn phases and mainly involved in cellular transport and metabolic processes (Figure 2I, Figure IIIC and Table II Supplement). Furthermore, 21% of the upregulated transcripts fitted to clusters 5 and 7, representing genes regulating morphogenesis and cell-cycle and whose expression levels decreased in infant stage (Figure 2I, Figure IIIC and Table II Supplement). Concordantly, 22 of the 84 D-DNM genes in our HLHS cohort were found in most clusters.

Together, these findings indicate that HLHS may result from alterations in specific gene programs critical during fetal ventricular cardiogenesis and suggest a role of cell-cycle and CM maturation as potential disease drivers.

### Specific gene networks are dynamically altered during differentiation of HLHS iPSCs towards CPs and CMs

To better dissect to which extent transcriptional alterations of native HLHS CMs underlie disease pathogenesis rather than physiological differences at the time of RV sampling, we generated iPSCs from 3 HLHS patients and 3 healthy subjects using Sendai-mediated reprogramming (Figures IV and V Supplement) and mechanistically analyzed developmental processes during early *in-vitro* cardiogenesis. Selection of HLHS patients was based on echocardiographic LV morphology (“thickened” phenotype) and presence of D-DNMs in genes affected twice or belonging to multihit gene families (Figure VI Supplement).

Control and HLHS iPSC lines were directed to early CPs^32^ and CMs^33^ using stepwise differentiation protocols (Figure 3A). We performed RNAseq at various time points and explored functional characteristics of HLHS DEGs by GSEA (Figure 3B and Table III Supplement). Dynamic alterations in several gene categories were common in both CP and CM differentiation protocols, including heart/aorta development, cell-cycle, and chromatin modification. Interestingly, unique aberrations in autophagy terms were present when directing HLHS iPSCs towards early CPs, while apoptosis-associated pathways appeared solely affected in later CPs (D6) and CMs (D8 and D14) (Figure 3B and Table III Supplement).

**Figure 3.**
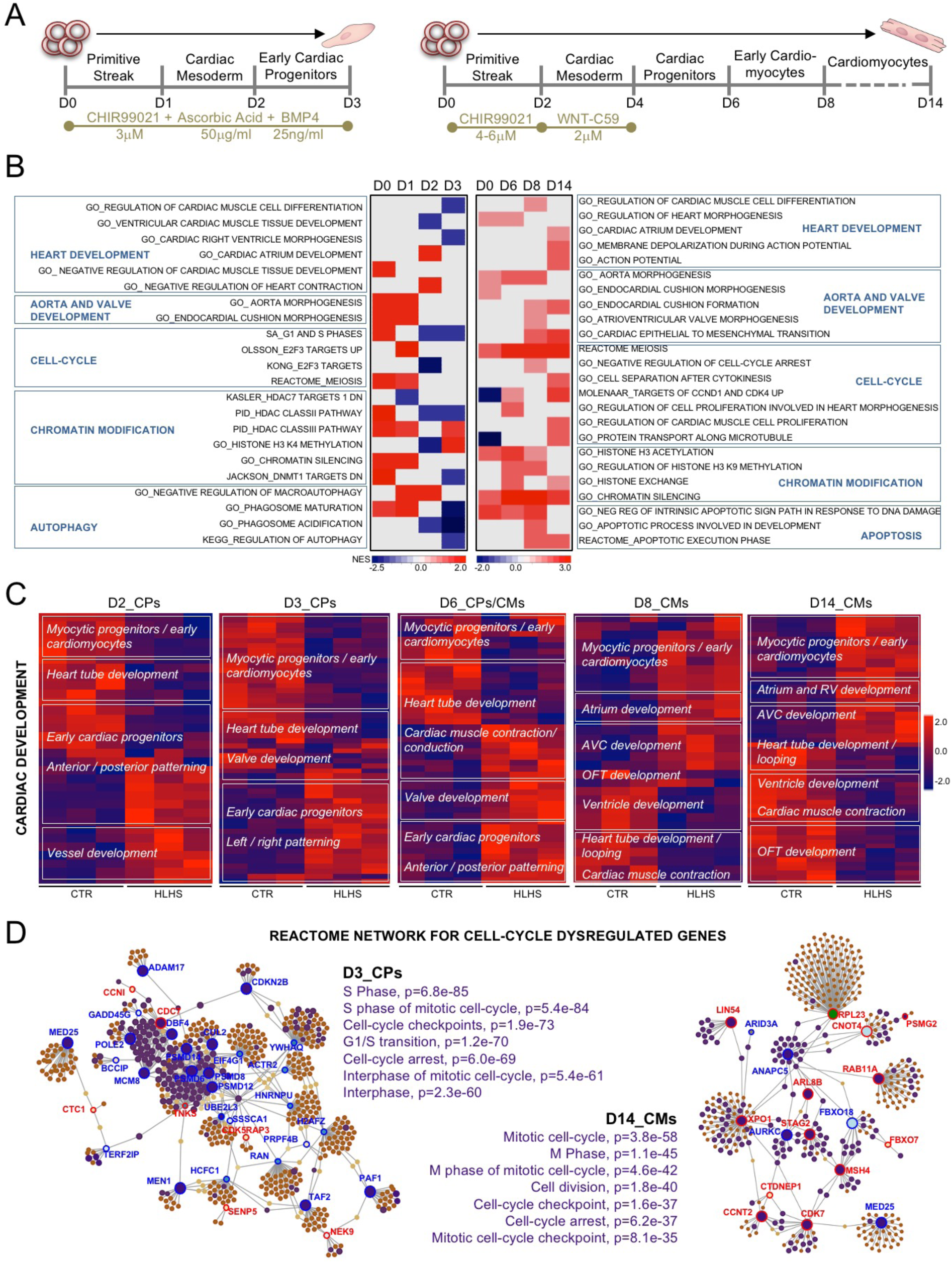
Gene expression analysis during iPSC-based cardiogenesis reveals networks of dysregulated genes in HLHS. **A**, Directed iPSC cardiac differentiation protocols used in the study. **B**, Heatmap of normalized enrichment scores (NES) for selected GSEA terms. Red and blue denotes terms with positive and negative NES, respectively. **C**, Heatmaps showing gene expression of DEGs (1.5-fold-expression, *p*-value ≤0.05) involved in cardiac development at the indicated days of differentiation. Values are row-scaled to show their relative expression. Blue and red are low and high levels respectively. **D**, Networkanalyst-generated protein-protein interactome of DEGs involved in cell-cycle at D3 and D14. Upregulated (red) and downregulated (blue) genes are shown. In purple are highlighted the genes belonging to the enriched GO categories specified on the side of the plots. Protein–protein interactions are indicated as solid gray lines between genes.

In CPs at D2 and D3, detailed analysis of DEGs involved in cardiac development revealed that genes expressed in committed myocytic precursors and important for heart tube formation (e.g. *ID2*, *TPM2*, *XIRP1*, *SRF, ETV1*) were downregulated in HLHS, while genes involved in anterior/posterior patterning (*HOXB9*) and in vessel/valve development (*VEGFB*, *TGFB2*, *GATA5*) were upregulated (Figure 3C). Transcripts typical of early CPs were decreased at D2, but augmented at D3, suggesting incomplete/delayed CP lineage specification. Concordantly, at D8 and D14, HLHS CMs showed upregulation of genes distinctive of myocytic progenitors/early immature CMs and altered expression of transcripts important for OFT and atrioventricular-canal (AVC) (*MEIS1*, *ISL1*, *TGFB2*, and *JUN*) as well as heart chamber development (*NR2F1*, *WNT2*, *ETV2*, *RXRA*) (Figure 3C and Table III Supplement), supporting dysregulated lineage-specific CM differentiation.

In CPs at D3, 74 of the 890 DEGs related to cell-cycle GO categories (Figure VIIA and Table III Supplement). Thirty-four of them generated a functional interactome network encompassing cell-cycle interphase pathways as the top enriched, with most leading genes being downregulated (Figure 3D and Table III Supplement). In CMs at D14, cell-cycle DEGs (42/754) generated 18 functional interaction nodes; top enriched terms within the interactome and regulation of the leading genes pointed to alteration in M phase, with active separation of chromatids (*STAG2*, *XPO1*) but defective progression through mitosis (*ANAPC5*) and cytokinesis (*AURKC*) (Figure 3D and Table III Supplement). Cell-cycle defects in HLHS CMs were also confirmed by proteomic analysis (Expanded Results, Figure VIII, and Table III Supplement).

Together, the specific transcriptional alterations detected during early CP specification and CM differentiation of HLHS iPSCs suggest the primary onset of the disease occurs at the initial stages of cardiogenesis when CM lineage decisions arise within CP populations.

### Defects in UPR-induced autophagy lead to delayed and disrupted CP lineage specification

To investigate the role of cell-cycle disturbances in HLHS pathogenesis, we analyzed cell-cycle patterns during early CP formation and compared them with the emergence of CP lineages marked by ISL1, NKX2-5, and TBX5. While control lines demonstrated G1 lengthening starting between D1 and D2, patient lines prolonged G1 phase with a 24h delay (Figure 4A). Concurrently, at D1, activation of *ISL1* and *NKX2-5* transcripts was dramatically reduced in HLHS (Figure VIIB Supplement) and correlated with significant lower proportions of cells expressing ISL1 and NKX2-5 proteins at D2 (Figure 4B), supportive of a retarded CP specification. Interestingly, four patterns of ISL1 and NKX2-5 expression were detected at D3: **i**) ISL1^low^/NKX2-5^low^, representing “early committed CPs”; **ii**) ISL1^high^/NKX2-5^high^, denoting “fully committed early CPs”; **iii**) ISL1^high^/NKX2-5^low^, typical of SHF progenitors; and **iv**) ISL1^low^/NKX2-5^high^, distinctive of FHF cells (Figure 4C). TBX5, a specific FHF marker, was mainly found in ISL1^low^/NKX2-5^high^ cells (Figure 4C), confirming FHF identity. The relative distribution of CP subgroups was altered in HLHS settings (Figure 4D). Importantly, all HLHS lines failed to upregulate *TBX5* during CP specification (Figure VIIB Supplement) and only few of the ISL1^low^/NKX2-5^high^ FHF cells expressed TBX5 (Figure 4E), indicating common defective transcriptional programs within the FHF lineage. To assess the contribution of apoptosis/proliferation to the observed CP phenotypes, we analyzed caspase-3 activation and EdU incorporation. Apoptosis was barely detectable in both HLHS and controls (Figure VIIC Supplement), arguing against any cell selection of HLHS CPs. Consistent with the alteration in cell-cycle patterns, global changes in cell proliferation rates between control and HLHS were only significantly different at D2 (Figure VIID Supplement). However, when analyzed separately at D3, ISL1^low^/NKX2-5^high^ FHF progenitors demonstrated higher proliferation in all HLHS lines (Figure 4F), consistent with the reported role of TBX5 as negative regulator of cell proliferation during early cardiac development^34^. Collectively, these data indicate common defects in CP lineage commitment and imbalance of both progenitor fields during initial steps of HLHS cardiogenesis.

**Figure 4.**
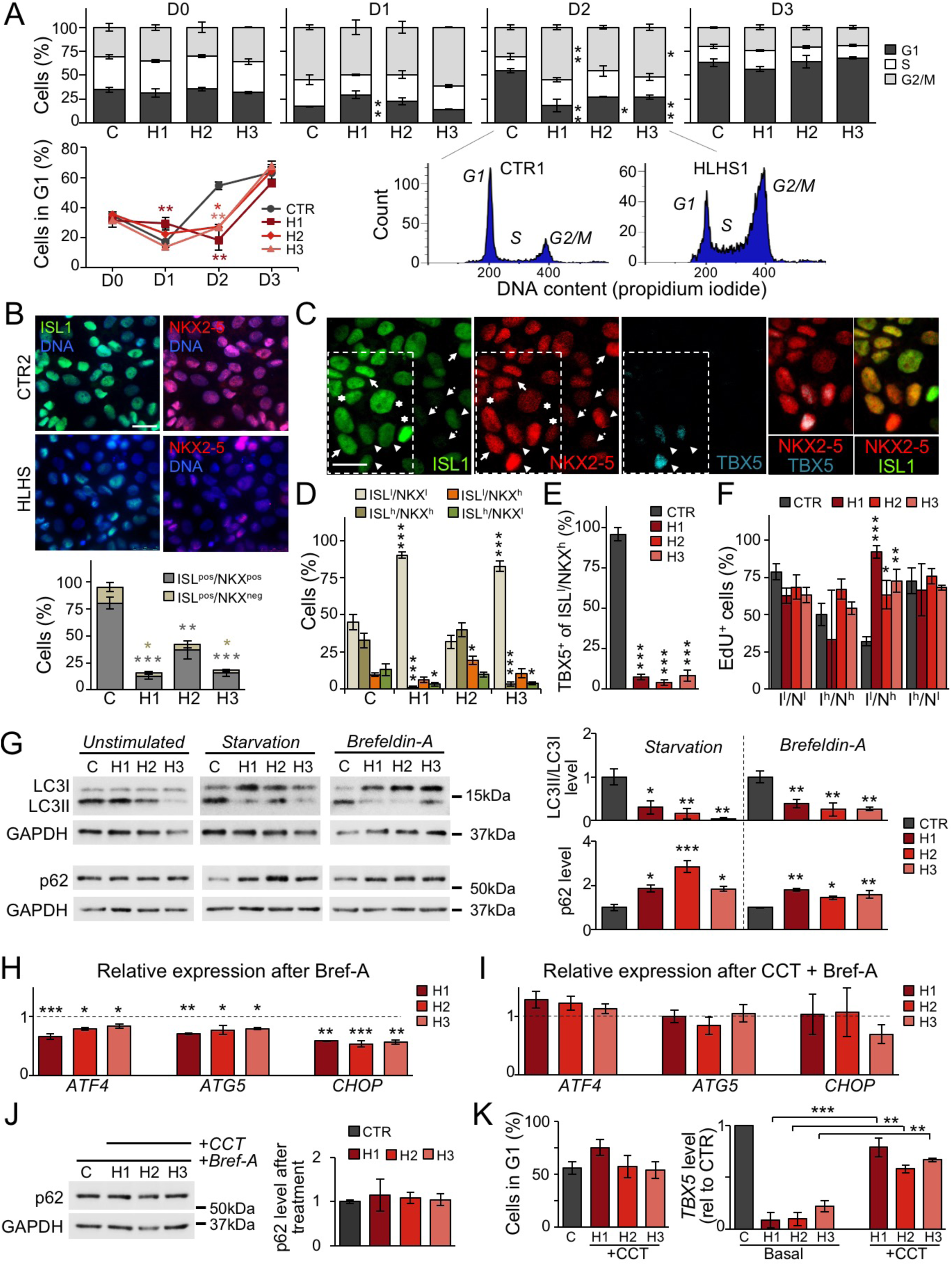
Defects in UPR and autophagy delay and disrupt CP lineage specification. To better discriminate possible phenotypic variations among diseased lines, functional results from each HLHS line are shown separately. Data from control lines have been combined. **A**, Propidium iodide staining analysis of HLHS (H) and control cells (C) during CP differentiation. Data are mean ± SEM, n=2-4 differentiations per line, N≥20000 cells per sample at each time point. *p<0.05, **p<0.01 compared to CTR (two-way ANOVA). **B**, Immunofluorescence analysis of ISL1 and NKX2-5 in HLHS and control CPs at D2. Scale bar, 25 μm. Data are mean ± SEM, 431 (CTR), 463 (HLHS1), 442 (HLHS2) and 396 (HLHS3) cells from n=3 differentiations per line. *p<0.05, **p<0.01, ***p<0.001 compared to CTR (one-way ANOVA). **C**, Representative immunofluorescence of ISL1, NKX2-5 and TBX5 in control CPs (CTR3) at D3. Four ISL1/NKX2-5 expression patterns are highlighted: ISL1^low^/NKX2-5^low^ (dotted arrows), ISL1^high^/NKX2-5^high^ (arrows), ISL1^high^/NKX2-5^low^ (asterisks), and ISL1^low^/NKX2-5^high^ (arrow heads). Scale bar, 20 μm. **D**, Distribution of cells with ISL1/NKX2-5 expression patterns from (C) in HLHS and control CPs at D3. Data are mean ± SEM, 369 (CTR), 347 (HLHS1), 322 (HLHS2), 357 (HLHS3) cells from 3 differentiations per line. *p<0.05, ***p<0.001 compared to CTR (one-way ANOVA). **E**, Percentage of ISL1^low^/NKX2-5^high^ cells expressing TBX5 in HLHS and control CPs at D3. Data are mean ± SEM, 114 (CTR), 70 (HLHS1), 182 (HLHS2) and 123 (HLHS3) cells from n=3 differentiations per line. ***p<0.001 compared to CTR (one-way ANOVA). **F**, Quantification of EdU^+^ cells in HLHS and control CP subpopulations at D3. Data are mean ± SEM, 369 (CTR), 347 (HLHS1), 322 (HLHS2), 357 (HLHS3) cells from 3 differentiations per line. *p<0.05, **p<0.005, ***p<0.001 compared to CTR (one-way ANOVA). **G**, Western blot of LC3 and p62 in HLHS and control CPs at D3 with and without starvation or brefeldin-A. For detection of LC3, all three conditions were carried out in presence of chloroquine. Data are mean ± SEM, n=2-3 differentiations per line. *p<0.05, **p<0.01, ***p<0.001 compared to CTR (one-way ANOVA). **H** and **I**, Expression analysis of *ATF4* and its downstream targets *ATG5* and *CHOP* in HLHS and control CPs at D3 after treatment with brefeldin-A alone (H) or in combination with the PERK activator CCT020312 (I). Shown are expression levels relative to controls. Data are mean ± SEM, n=2-4 differentiations per line. *p<0.05, **p<0.01, ***p<0.001 compared to CTR (one-way ANOVA). **J**, Western blot of p62 in HLHS and control CPs at D3 after treatment with brefeldin-A and CCT020312. Data are mean ± SEM, n=3-6 differentiations per line. **K**, Propidium iodide staining-based quantification of cells in G1 phase in HLHS and control CPs at D2 (left) and *TBX5* expression by qRT-PCR at D3 (right) after 6h-treatment of HLHS cells with CCT020312 at D1.5. Data are mean ± SEM, n=2-3 differentiations per line, N≥20000 cells per sample in left panel. **p<0.01, ***p<0.001 compared to own basal (one-way ANOVA).

Autophagy and cell-cycle are coordinated and reciprocally regulated^35^. Since autophagy was altered in HLHS CPs from D1 on (Figure 3B), we measured autophagic flux in cells at D3 by analyzing the levels of LC3II, a marker of autophagosomes, and p62, a substrate for autophagic degradation. Under basal conditions, LC3II and p62 proteins were normal in all HLHS lines (Figure 4G and Figure VIIE Supplement). However, after activation of autophagy (starvation or brefeldin-A), defective autophagosome formation and p62 degradation were evident in all HLHS lines (Figure 4G). This was also confirmed by Cyto-ID staining of autophagic vacuoles (Figure VIIF Supplement). Brefeldin-A triggers autophagy *via* ER stress and activation of the unfolded protein response (UPR)^36^, a pathway also challenged by starvation. UPR plays an important role in cell fate acquisition of embryonic stem cells and progenitors^37–39^. Given the recovery of HLHS D-DNMs in ER-stress/UPR genes (Figure 1D and Table I Supplement), we asked whether the defective autophagy in HLHS CPs is caused by impaired UPR. Upon ER stress, UPR is transduced by de-repression of three ER membrane proteins: IRE1α, PERK, and ATF6 that work alone or in concert to restore normal cellular function^40^. Interestingly, neither an increased splicing of *XBP1* mRNA, which occurs downstream of IRE1α stimulation, nor activation of *ATF6* were altered in HLHS CPs at D3 upon brefeldin-A treatment (Figure VIIG Supplement). Instead, we measured a specific common defect of all HLHS lines in activating the PERK pathway, as indicated by decreased PERK-mediated phosphorylation of eIF2α (Figure VIIH Supplement) and reduced increase of *ATF4* and its downstream targets including *CHOP* (Figure 4H and Figure VIIG Supplement). A six-hour treatment of HLHS cells with the selective PERK activator CCT020312 normalized *ATF4* and *CHOP* levels and rescued the defective activation of autophagy (Figures 4I and 4J). Importantly, early application of CCT020312 at D1.5 of CP differentiation was sufficient to revert the HLHS phenotype, as indicated by normalization of the number of cells in G1 at D2 and the level of *TBX5* at D3 (Figure 4K). Conversely, HLHS-like disturbances in G1-phase lengthening and *TBX5* upregulation could be induced in control cells by inhibiting autophagy at D1.5 using chloroquine, with no influence on *ISL1* expression (Figure VIII Supplement).

Taken together, these results suggest that defects in UPR/autophagy activation in the early phase of CP specification contribute to delayed and disrupted CP lineage commitment in HLHS.

### Single-cell RNAseq defines impaired CM lineage segregation and maturation in HLHS

To investigate the consequences of altered CP specification on CM-subtype formation, we performed single-cell RNAseq of early CMs at D14. Transcriptomes of 6,431 control and 4,439 HLHS cells were recovered. Unsupervised clustering analysis identified 10 distinct sub-populations (clusters 0-9) (Figure 5A and Table III Supplement). We assigned identities to each population by cross-referencing the most highly and uniquely expressed genes in each cluster with known cardiac subtype markers from human and mouse single-cell studies (Figure 5B and Expanded Results Supplement)^28, 29, 41–43^. We captured transcriptome characteristics of primary (OFT and AVC) and chamber myocardium, early and late CPs, maturing CMs, and proliferating CMs in G1/S and G2/M phases (Figure 5A through 5C and Figure IXA Supplement). Remarkably, HLHS cells contributed almost exclusively to CP clusters (cluster 7: CTR=40, HLHS=450; cluster 9: CTR=35, HLHS=117) and were strongly under-represented in the cluster containing terminally differentiated CMs (cluster 8: CTR=318, HLHS=37), together suggestive of an intrinsic differentiation delay. Moreover, we noticed a significant reduced proportion of HLHS cells in the proliferating cell clusters (cluster 4: CTR=922, HLHS=284; cluster 6: CTR=675, HLHS=103), corroborating a premature cell-cycle exit in HLHS CMs. Interestingly, co-expression of LV enriched genes was observed mainly in CMs of cluster 0 that was scarce in HLHS cells (cluster 0: CTR=1446, HLHS=497). This correlated with an overall downregulation of LV transcripts in HLHS (e.g. *TBX5*, *HAND1*, *SLIT2*, *GJA1*) (Figure IXB Supplement), hinting to a potentially reduced LV commitment. Differential expression analysis within each cluster revealed increased levels of pro-apoptotic genes (*TNFRSF12A*, *FAM162A*) and reduced expression of anti-apoptotic regulators (e.g. *MTRNR2L1*) in HLHS CMs (Figure 5D and Figure IXC Supplement), confirming bulk RNAseq results. Moreover, HLHS CMs presented an overall upregulation of genes involved in glucose catabolism, downregulation of mitochondrial transcripts, and lower expression of sarcomeric genes; exceptions were *MYL7* and *TNNI1*, isoforms highly expressed in immature CMs (Figure 5D and Figure IXC and IXD Supplement), collectively suggesting reduced cellular differentiation/maturation. To further verify this, we used recently published single-cell profiles of healthy iPSC-CMs at different stages of differentiation^44^. We ordered CMs captured at D5 and D14 together with our cells in pseudotime. The resulting differentiation trajectory began with CM at D5 and bifurcated into two lineages where CMs at D14 were allocated (Figure 5E). Importantly, a large proportion of HLHS CMs, although at D14, clustered with D5 CMs and expressed high level of *KRT19*, a gene specific of early immature CMs (Figure 5E).

**Figure 5.**
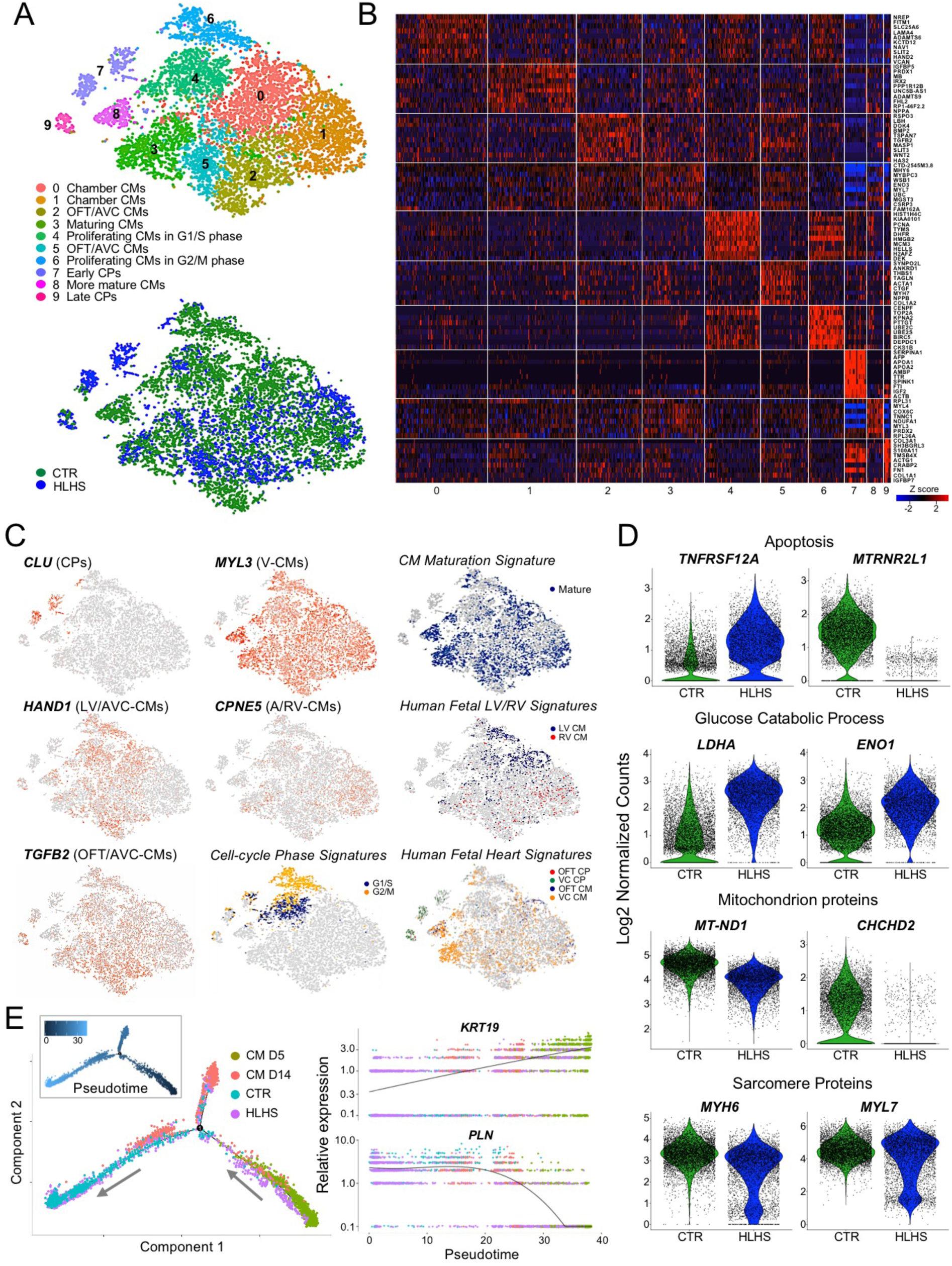
Single-cell RNAseq of iPSC-CMs reveals defects in cardiac lineage segregation and maturation in HLHS. **A**, t-SNE plot of all HLHS and CTR cell populations captured at D14 colored by cluster identity (top) and genotype (bottom). Data are from 2 control (CTR2 and CTR3) and 2 HLHS (HLHS2 and HLHS3) lines. **B**, Heatmap showing Z score scaled average expression levels of the top ten DEGs for each cellular cluster. **C**, Expression of selected genes marking subpopulations on t-SNE plot. Single gene panels: red and gray indicates high and low expression, respectively. Signature panels: color key indicates cells matching with the gene signatures tested (see Expanded Methods). A, atria; VC, ventricular chamber. **D**, Violin plots of selected DEGs between CTR and HLHS cells. All genes represented have a *p*-value <0.05. **E**, Branching analysis of HLHS and CTR CMs at D14 together with CMs at D5 and D14 from^44^ colored by genotype and estimated pseudotime along the inferred cell trajectory (inset). Pseudotime dynamics of early (*KRT19*) and mature (*PLN*) CM genes in dependence on inferred cell identity.

Together, these results indicate that both early CM-subtype lineage specification and CM differentiation/maturation are disrupted in HLHS. Moreover, they confirm, at a single-cell transcriptional level, a premature cell-cycle withdrawal in diseased CMs.

### 3D heart patches unravel dysregulated nodes for CM contractility, maturation, and survival in HLHS

We further analyzed HLHS phenotypes in a multicellular 3D context by repopulating decellularized scaffolds from non-human primate LV heart slices with early D14 iPSC-CMs. Standardized cell seeding was achieved using bioprinting and constructs underwent electromechanical conditioning (1Hz pacing, 1mN diastolic preload) in customized biomimetic chambers^45^ for 24 days (Figure 6A). Contractile force of HLHS patches was significantly reduced compared to controls and did not increase, but rather decreased, overtime (Figure 6B and 6C). Moreover, while progressive electromechanical maturation was evident in controls, HLHS tissues failed to respond to high stimulation frequencies and to develop a positive force-frequency relationship (Figure 6D through 6F). Concurrently, Ca^2+^ imaging of single CMs within the patches demonstrated a gradual decline in the number of electrically responsive HLHS cells (Figure 6G through 6I) and intrinsic Ca^2+^ handling defects, with failed increase of systolic and abnormal elevation of diastolic Ca^2+^ at high stimulation rates (Figure 6J and 6K). Expression profile of HLHS CMs isolated from 3D patches at D12 confirmed inherent molecular changes of pivotal genes involved in electromechanical coupling and Ca^2+^ homeostasis (Figure 6L). Moreover, live evaluation of cell viability revealed a progressive increase of dead cells in HLHS tissues (Figure XA Supplement), suggesting apoptosis as additional mechanism underlying the functional deterioration. Immunohistochemistry demonstrated a gradual increase of apoptotic CMs (clCasp3^+^/TUNEL^+^) in diseased patches (Figure 7A). Intriguingly, not all TUNEL^+^ CMs expressed activated caspase-3, while the opposite was true; moreover, the percentage of clCasp3^-^/TUNEL^+^ cells was also significantly elevated in HLHS (Figure 7A), indicating apoptosis-independent DNA damage. To better assess this, we performed co-immunofluorescence analysis of activated caspase-3 and activated p53 (phP53), a tumor suppressor that induces either cell-cycle arrest to facilitate DNA repair or apoptosis^46^. We detected clCasp3^+^/phP53^+^ and clCasp3^-^/phP53^+^ CMs, with a significant increase of both in HLHS tissues (Figure 7B). Notably, the percentage of clCasp3^-^/phP53^+^ cells matched well with the proportion of clCasp3^-^/TUNEL^+^ cells, suggesting that DNA damage might be the first event leading to apoptosis of HLHS CMs.

**Figure 6.**
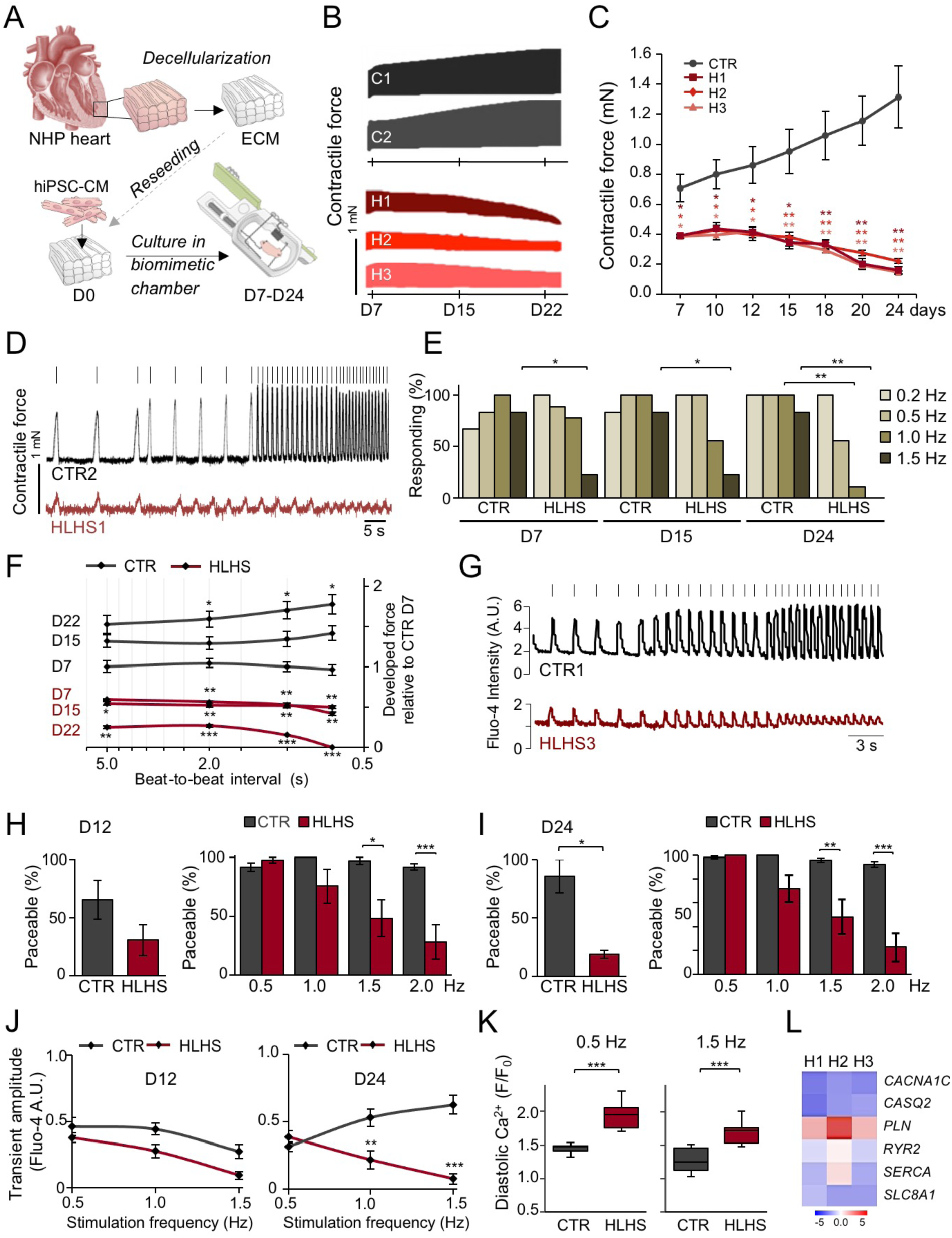
Three-dimensional culture of iPSC-CMs under electromechanical stress reveals HLHS-related functional abnormalities. **A**, Schematic of the experimental setup for 3D culture of iPSC-CMs within decellularized heart patches kept in biomimetic chambers providing mechanical load and electrical stimulation while allowing continuous monitoring of force development. All measurements were done in patches generated from 2 control and 3 HLHS lines. Unless otherwise illustrated, results from different control and HLHS lines have been pooled. NHP, non-human primate. **B** and **C**, Representative plots (B) and statistical analysis (C) of contractile force in HLHS and control patches over 24 days of culture. In (C), data are mean ± SEM of serial measurements at the indicated days, n=8 (CTR), n=3 (HLHS1 and HLHS2), n=6 (HLHS3) patches. *p<0.05, **p<0.01 compared to CTR (two-way repeated-measures ANOVA). **D**, Representative traces of contraction force at increasing stimulation frequencies in one control and one HLHS line from an experiment aimed at assessing the force-frequency relationship and the paceability at different stimulation frequencies. Stimulation pulses are indicated as vertical bars above the respective tracing. **E**, Percentage of patches responding to stimulation at indicated pacing frequencies. n=6 (CTR) and n=9 (HLHS). *p<0.05, **p<0.01 compared to CTR at the same day and frequency (Fisher’s exact test). **F**, Force-frequency relationship (FFR), depicted as the developed force (normalized to the mean force developed by CTR patches at day 7) as a function of the beat-to-beat interval (depicted on a logarithmic scale). Serial FFR values obtained from HLHS (n=11) and control (n=8) patches at indicated time points are shown. Data are mean ± SEM. *p<0.05, **p<0.01, ***p<0.001 compared to CTR at D7 at the same beat-to-beat interval (mixed effects model). **G**, Representative images of Fluo-4-based intracellular calcium transients from single control and HLHS CMs within the patch at increasing pacing rates. Vertical bars over the tracings represent the stimulation pulses. **H** and **I**, The left graph shows the overall percentage of paceable CMs within control and HLHS patches at D12 (H) and D24 (I) of 3D culture. The right graph shows, only considering cells that were paceable with at least one of the applied pacing rates, the percentage of cells responding at the indicated frequencies at D12 (H) and D24 (I). Data are mean ± SEM, n=6 (CTR) and n=9 (HLHS) patches; left panels: N=183 and 156 (CTR), N=466 and 70 (HLHS) cells in (H) and (I), respectively; right panels: N≥67 and 90 (CTR), N≥21 and 17 (HLHS) cells for each frequency in (H) and (I), respectively. *p<0.05, **p<0.01, ***p<0.001 compared to CTR (Mann Whitney test for left panels and two-way ANOVA for right panels). **J**, Amplitude of the systolic calcium transients plotted as a function of the stimulation frequency for CMs within control (n=6) and HLHS (n=9) patches at D12 and D24 of 3D culture. N≥65 and 81 (CTR), N≥3 and 35 (HLHS) cells for each frequency at D12 and D24, respectively. Data are mean ± SEM. **p<0.01, ***p<0.001 compared to CTR (two-way ANOVA). **K**, Diastolic calcium level (expressed as ratio of the diastolic Fluo-4 intensity at the indicated pacing frequency (*F*) and the basal Fluo-4 intensity at the beginning of the experiment (*F_0_*)) of single CMs in D24 control (n=6) and HLHS (n=9) patches at 0.5 Hz and 1.5 Hz pacing rates. N=91 and 83 (CTR), N=70 and 44 (HLHS) cells at 0.5 Hz and 1.5 Hz, respectively. Data are mean ± SEM. ***p<0.001 (Mann Whitney test). **L**, Expression level of key genes for electromechanical coupling (black) and Ca^2+^ homeostasis (red) in CMs isolated from HLHS and control 3D patches at D12. Data are log2 mean fold changes relative to controls, n=3 patches per line.

**Figure 7.**
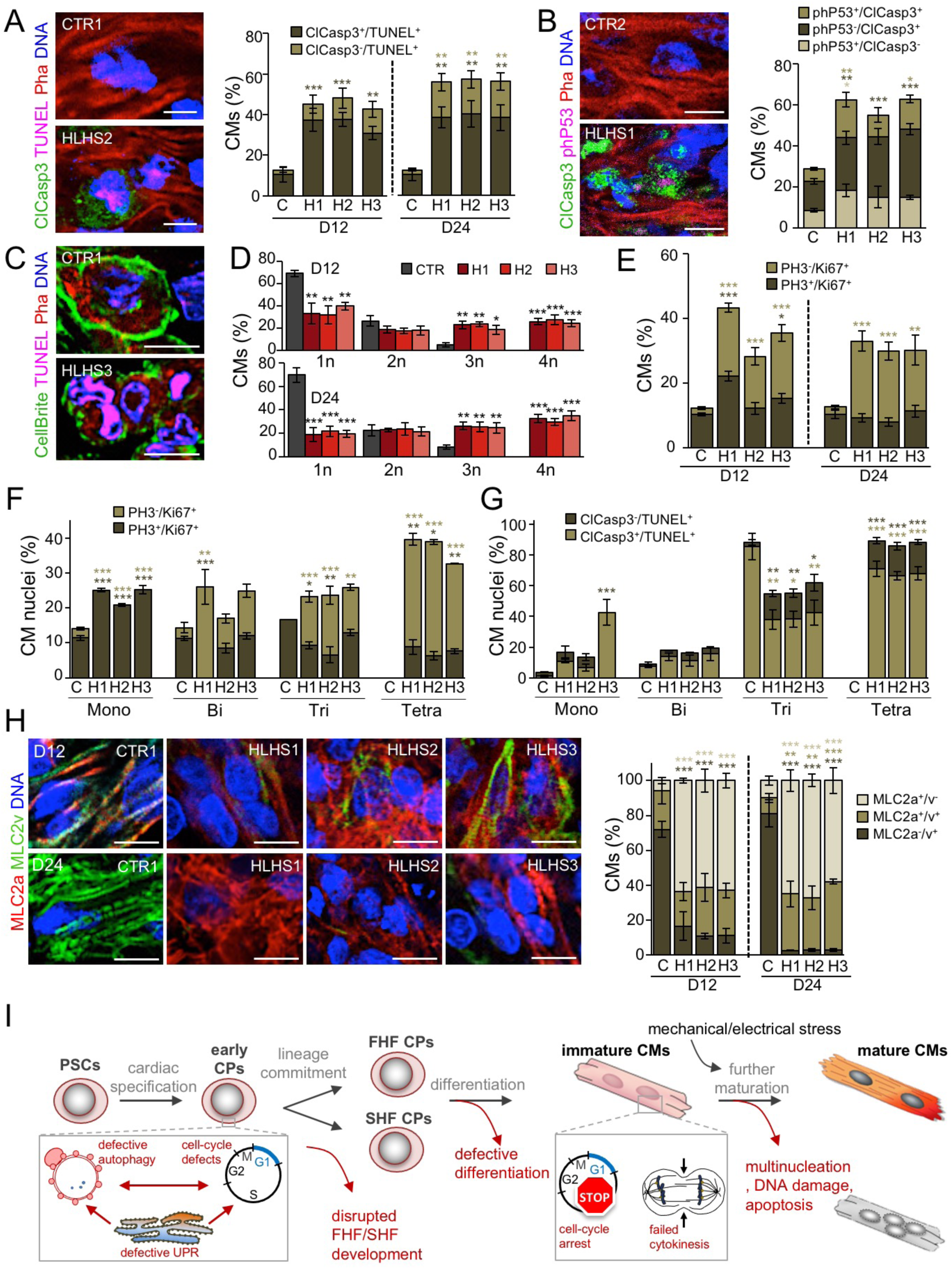
Aberrant apoptosis, multinucleation, and maturation of HLHS CMs in 3D biomimetic culture. All measurements were done in patches generated from 2 control and 3 HLHS lines. Results from the 2 different controls have been pooled. **A**, Representative fluorescence images of D24 control and HLHS patches after immunostaining for activated caspase 3 (ClCasp3) in conjunction with TUNEL labeling. Phalloidin (Pha) marks F-actin and distinguishes CMs. Scale bar, 10 µm. Bar graph shows the statistical evaluation of D12 and D24. Data are mean ± SEM, n=8 (CTR, N=226 cells) and n=4 (each HLHS line, N≥200 cells per line) patches at D12; n=7 (CTR, N=534) and n=3 (each HLHS line, N≥434 cells per line) patches at D24. **p<0.01, ***p<0.001 compared to CTR (one-way ANOVA). **B**, Representative immunostaining for activated Caspase 3 (ClCasp3) and phosphorylated P53 (phP53) in D24 control and HLHS patches. Scale bar, 10 µm. Bar graph illustrates the percentages of CMs (identified by Pha) expressing one or both markers. Data are mean ± SEM, n=10 (CTR, N=652 cells), n=6 (HLHS1, N=572 cells), n=5 (HLHS2, N=470 cells), and n=5 (HLHS3, N=314 cells) patches. *p<0.05, **p<0.01, ***p<0.001 compared to CTR (one-way ANOVA). **C**, Representative fluorescence images of CMs (identified by Pha) after plasmamembrane (CellBrite), TUNEL, and DNA labeling within control and HLHS patches at D24. Scale bar, 10 µm. **D**, Bar graphs showing the percentage of CMs with one (1n), two (2n), three (3n) and four (4n) nuclei in the different cell lines at D12 and D24. Data are mean ± SEM, n=6 (CTR, N=340 cells) and n=3 (each HLHS line, N≥182 cells per line) patches at D12; n=6 (CTR, N=341) and n=3 (each HLHS line, N≥147 cells per line) patches at D24. *p<0.05, **p<0.01, ***p<0.001 compared to CTR (one-way ANOVA). **E**, Bar graphs showing the percentage of CMs positive for phosphorylated histone 3 (PH3) and Ki67 at D12 and D24. Data are mean ± SEM, n=7 (CTR, N=528 cells), and n=7 (HLHS1, N=698 cells), n=5 (HLHS2, N=463), and n=5 (HLHS3, N=668) patches at D12; n=7 (CTR, N=673 cells), and n=7 (HLHS1, N=682 cells), n=5 (HLHS2, N=486), and n=5 (HLHS3, N=485) patches at D24. *p<0.05, **p<0.01, ***p<0.001 compared to CTR (one-way ANOVA). **F,** Bar graph showing the percentage of nuclei positive for PH3 and Ki67 in mono-, bi-, tri-, and tetra-nucleated CMs at D24. Data are mean ± SEM, n=3 patches per line; N=331 (CTR), N=258 (HLHS1), N=228 (HLHS2), and N=240 (HLHS3) cells. *p<0.05, **p<0.01, ***p<0.001 compared to CTR (one-way ANOVA). **G**, Bar graph showing the percentage of nuclei positive for ClCasp3 and phP53 in mono-, bi-, tri-, and tetra-nucleated CMs at D24. Data are mean ± SEM, Data are mean ± SEM, n=7 (CTR) and n=3 (each HLHS line) patches; N=401 (CTR), N=194 (HLHS1), N=175 (HLHS2), and N=237 (HLHS3) cells. *p<0.05, **p<0.01, ***p<0.001 compared to CTR (one-way ANOVA). **H**, Left, representative immunostains for MLC2a and MLC2v in control and HLHS patches. Scale bar, 10 µm. Right, bar graph shows statistical evaluation. Data are mean ± SEM, n=8 (CTR, N=951 cells) and n=4 (each HLHS line, N≥428 cells per line) patches at D12; n=8 (CTR, N=949) and n=4 (each HLHS line, N≥442 cells per line) patches at D24. **p<0.01, ***p<0.001 compared to CTR (one-way ANOVA). **I**, Scheme depicting the identified steps during cardiac development at which HLHS-related abnormalities interfere with normal development and contribute to the complex CHD phenotype.

An interesting finding emerging from histological analyses was abnormal multinucleation of CMs in HLHS patches, already evident at D12 (Figure 7C). Polyploidy and binucleation are characteristic features of mammalian CMs that develop shortly after birth when most differentiated CMs exit cell-cycle^31^. In humans, polyploidy is often increased in pathological conditions; however, binucleation occurs in only 25% CMs, with no evidence of tri- or tetranucleation^47^. In control patches, most CMs carried 1 (∼70%) or 2 (∼25%) nuclei, and only few cells presented 3 or more; conversely, in HLHS tissues, ∼50% of CMs were tri/tetranucleated (Figures 7C and 7D), indicating an intrinsic failure to complete cytokinesis and premature cell-cycle withdrawal. Markers of cell-cycle activity (Ki67) and mitosis (phospho-histone H3, PH3) demonstrated an overall increase in the number of active CMs in HLHS patches, with the proportion of cells in mitosis being similar (Figure 7E). However, detailed analysis of Ki67 and PH3 expression revealed that, in mononucleated CMs, most nuclei were in M phase (Ki67^+^/PH3^+^) and their percentage was higher in HLHS (Figure 7F). Importantly, greater the degree of multinucleation in HLHS, higher was the proportion of Ki67^+^/PH3^-^ *vs* Ki67^+^/PH3^+^ nuclei (Figure 7F), suggesting that polyploidization was also occurring in multinucleated diseased CMs. To assess whether and how the abortive cell-cycle mode of HLHS CMs related to apoptosis, we evaluated the distribution of TUNEL^+^ nuclei and clCasp3^+^/TUNEL^+^ cells (Figure 7G). We found a striking correlation between the number of TUNEL^+^ nuclei and the level of multinucleation, being most damaged nuclei in the tri-/tetranucleated CMs. The latter showed also the highest percentage of activated caspase-3 (Figure 7G), together indicating chromosomal instability acquired during aberrant cell-cycle as possible apoptotic trigger in HLHS CMs.

We hypothesized that the observed HLHS CM phenotype might be a specific reaction to strong cues for cell division/growth and maturation, which are provided by the electromechanical preconditioning in the 3D-tissue environment. Since our transcriptome profiling of early HLHS CMs developed in 2D-monolayer pointed to defects in differentiation/maturation, we performed co-immunofluorescence analysis for MLC2a (expressed in all immature CMs and becoming atrial-specific after terminal chamber differentiation) and MLC2v (expressed in maturing vCMs). In HLHS patches, we measured a striking increase of MLC2a^+^ cells with reduction of MLC2v^+^ CMs at D12 and D24 (Figure 7H); moreover, sarcomere organization was impaired (Figure XB Supplement), indicative of reduced maturation. Similar results were obtained in HLHS CMs (D30 and D60) in 2D-monolayers (Figure XC through XE Supplement). Expression analysis of terminally differentiated atrial, ventricular, and CM maturation markers confirmed that the abnormal distribution of HLHS MLC2a^+^/MLC2v^+^ cells was a result of defective ventricular differentiation/maturation rather than an atrial lineage switch (Figure XF Supplement). This was corroborated by measuring intracellular Ca^2+^ cycling of 2D CMs (Figure XIA through XIC Supplement). Importantly, no aberrant multinucleation or apoptosis were visible in 2D (Figure XID through XIF Supplement).

Together, these results indicate that impaired maturation and inability to respond to cues for developmental growth by normal progression through cell-cycle leads to increased apoptosis of vCMs contributing to HLHS pathogenesis.

## DISCUSSION

Our work provides a multidisciplinary framework for studying human heart development and its disruption in CHD. WES of HLHS parent-offspring trios and transcriptomics of patient RV CMs identified consistent perturbations in gene expression programs and associated features of abnormal cardiac development. Human iPSC lines derived from HLHS patients facilitated dynamic evaluation of transcriptional and cellular phenotypes during progression of cardiogenesis in single cells and 3D functional modeling of ventricular chamber development. Together, our data indicate that initial perturbations of the cell-cycle/UPR/autophagy hub result in disrupted differentiation of early CP lineages and disproportionate allocation of vCM-subtypes in HLHS. Moreover, impaired maturation and premature cell-cycle exit of vCM reduce their ability to respond to cues for tissue growth resulting in increased apoptosis and ventricular hypoplasia (Figure 7I).

### Identification of dysregulated transcriptional nodes in HLHS

Pathway analysis of D-DNMs in HLHS cases showed significant enrichment in heart development and its related signaling pathways. Comprehensive investigation of single-cell expression profile and dynamically regulated gene programs in CMs indicated an occurrence of D-DNMs in gene modules typical of the embryonic stage encompassing cell-cycle. Transcriptomes of infant HLHS vCMs closely resembled a fetal stage, demonstrating that embryonic gene programs persist in HLHS CMs after birth. We identified dysregulated transcriptional nodes common to both CP and CM states or unique of one state. Moreover, single-cell RNAseq of HLHS CMs recognized distinct regulatory defects in specific cellular subsets indicating that both early CM-subtype lineage specification, maturation and cell-cycle are disrupted in the disease.

Previous studies in HLHS iPSCs showed reduced cardiac differentiation and structural CM maturation in presence of dysfunctional NOTCH signaling^17–19^ and suggested defective commitment to the ventricular lineage^21^, corroborating our results. Moreover, abnormalities in endocardial cells were shown to influence CM proliferation/maturation in HLHS^48^.

Evidence of cell-cycle arrest and impaired maturation has been reported in vCMs of the hypoplastic LV of HLHS fetuses^49^ and *Ohia* mutants^10^. Loss of replication potential during fetal growth likely impacts cardiac chamber development and function. Indeed, our 3D functional studies revealed, both on the tissue and single CM level, abnormalities of excitation-contraction coupling reminiscent of failing human myocardium. Beside the valvular perturbations, chamber-specific differences in proliferation rates at early stages of development may explain why loss of CM proliferation in HLHS affects LV more than RV. By analyzing cell-cycle gene signature of single CMs from RV and LV regions of human embryonic hearts from 5 to 25 weeks of gestation^29^, we found LV CMs proliferate more than their RV counterpart at 5 weeks (58% LV *vs* 48% RV), but these differences extinguish later in development (45% and 55% LV *vs* 43% and 53% RV at 6 and 7 weeks, respectively). Concordantly, similar analysis using our single-cell RNAseq data of D14 control iPSC-CMs with LV and RV transcriptome revealed a ∼3-fold increased proliferation of LV compared to RV cells (34.5% LV *vs* 11.8% RV, p<0.05).

### Linking clinical phenotype to mechanism

HLHS is a spectrum of disease that includes LV and aortic hypoplasia with aortic and mitral valve malformations ranging from stenosis to complete atresia. Historically, the “flow-volume hypoplasia” hypothesis has been supported by experiments in multiple model systems. However, recent observations in patients^8, 9^ argue that hemodynamic conditions alone are insufficient to explain variable LV morphology. We focused on the “thickened” LV morphology with a clearly visible LV lumen^8, 9^ and were able to identify a common mechanism (Figure 7I) for the initial arrested LV development in this specific anatomic subtype. Intrauterine, fetal valvuloplasty in HLHS patients is being studied as treatment option to enable LV growth. However, the existence of intrinsic CM defects such as loss of replication and increased apoptosis in distinct subsets of HLHS patients suggest a way to potentially improve the unpredictable results of this aortic valvuloplasty approach^50^. Furthermore, knowledge of these specific defects allows development of rational strategies, either by stratification or adjunctive therapies, potentially permitting biventricular repair in those with less serious forms of LV hypoplasia.

### Limitation of the study

The small number of our HLHS parent-off-spring trios limited identification of new disease candidate genes. Regardless, the combination of WES data from our and the PCGC HLHS cohorts identified disturbed genetic pathways with significant burden enrichment in D-DNMs, which we functionally validated.

The inability to obtain LV samples from HLHS patients, differences in RV *vs* LV physiology, age, medications, and the intrinsic variability among different iPSC lines are potential confounders. However, the recovery of consistent molecular and cellular alterations in native RV CMs and *in-vitro* iPSC models reinforces their causal relation to HLHS pathogenesis.

Taken together, our results suggest that a shared mechanism for subsets of HLHS. Moreover, they highlight that reduced LV growth in HLHS is likely not the sole consequence of disrupted valve formation and impaired blood flow; instead, intrinsic defects of vCM lineage development are primary contributors to the disease pathogenesis. More broadly, our work illustrates that, despite the extensive genetic heterogeneity underlying CHD, studying cardiac developmental processes in CHD patients using converging multidimensional technologies can provide deep mechanistic insight into these complex diseases to suggest novel therapeutic approaches.

## ACKNOWLEDGMENTS

We thank the HLHS family members from the US and Europe and the healthy volunteers for participation in our study. We would like to thank A.M., M.K., and P.J.G. lab members for technical assistance, helpful suggestions, and discussions. We would like to acknowledge Birgit Campbell and Christina Scherb for their technical assistance in cell culture, Gabi Lederer (Cytogenetic Department, TUM) for karyotyping, members of the KaBi DHM Biobank for patient sample collection, Alma Kuechler (Institute of Human Genetics, University Hospital Essen) for providing one HLHS trio, and Rabea Hinkel (Deutsches Primatenzentrum Göttingen) for providing NHP heart samples.

## SUPPLEMENTAL MATERIAL

Expanded Methods

Expanded Results

Supplemental Figures I-XI

Supplemental Excel Tables I-IV

Supplemental Tables V-VI

References 51-75

