## Supplement for "Sequential defects in cardiac lineage commitment and maturation cause hypoplastic left heart syndrome"

### SUPPLEMENTAL MATERIAL

#### SUPPLEMENTAL METHODS

##### HLHS parent-offspring trios

Blood samples for WES were obtained from 87 probands diagnosed with a sporadic form of HLHS and their respective parents. The parent-offspring trios were recruited in Munich (37 trios), Utah (46 trios), Amsterdam (3 trios), and Essen (1 trio). From a subset of the patients, skin fibroblasts suitable for iPSC generation were obtained under the regulations of the cardiovascular biobank of the German Heart Center Munich (KaBi-DHM). Informed consent was obtained from all donors or their legal representatives. The study was approved by the local ethics committees at the respective institutions (KaBi-DHM: 5943/13; IHG: 5360/13). Additionally, *de-novo* variants recovered in HLHS patient-offspring trios from the Pediatric Cardiac Genomics Consortium (PCGC)[27] were used for complementary analyses.

##### Human heart samples

Human heart ventricular samples at three developmental stages (fetal (16-23 post-conception weeks); early infancy (2-10 months); adulthood (21-50 years)) as well as samples from HLHS patients (5-11 days) were obtained either from aborted fetuses, pathology or during the Norwood Stage I palliation. Ventricular samples in the surgical HLHS group were harvested from the free right ventricular free wall during establishment of the right ventricle-pulmonary artery (RV-PA) shunt. Ventricular samples from pathology were also harvested from the free right ventricular free wall to avoid differences based on localization. All samples were immediately snap frozen and stored in liquid nitrogen until use. For further procedures all samples were treated equally. Informed consent was obtained from the subjects or their legal representatives; the tissue collection was approved by the local ethics committee. The active Institutional Review Board- (IRB) approved protocol for the fetal samples is HSM 15-00696. The ethical vote for the forensic samples is 247/16S. The ethical vote for the material obtained during clinically-indicated cardiac surgical procedures is 5943/13 (KaBi-DHM).

##### iPSC line generation, cell culture, and cardiac differentiation

The iPSC lines were generated from 3 HLHS patients and 3 healthy subjects by reprogramming dermal fibroblasts or peripheral blood mononuclear cells using the CytoTune-iPS$^{TM}$-iPS 2.0 Sendai Reprogramming Kit (Thermo Fisher Scientific), as previously described[51,52]. Several iPSC clones from each HLHS patient and healthy control probands were manually picked and expanded for characterization. Sendai virus detection in iPSCs was performed by RT-PCR at passage 5 to 10. Expression of pluripotency markers in iPSCs was verified by RT-PCR and immunostaining. Spontaneous differentiation of iPSCs into cells of all three germ layers was induced by embryoid body (EB) formation, as previously described[53]. Expression of lineage markers representative for three embryonic germ layers was assessed at day 21 of EB differentiation. Karyotyping and testing for karyotype-specific anomalies was done for all iPSC lines.

iPSCs were maintained in TeSR$^{TM}$-E8$^{TM}$ medium on matrigel-coated dishes and non-enzymatically passaged every 5–7 days after reaching 85–100% confluency using 0.5 mM EDTA/PBS solution.

For directed cardiac progenitor (CP) differentiation, iPSCs were collected using Accutase (StemCell$^{TM}$ Technologies) and plated on geltrex-coated dishes at a density of $5 \times 10^4$ cells/cm$^2$ in CP-induction medium, containing DMEM/F-12, 1x B-27 Supplement without vitamin A, 2 mM L-glutamine, 100 U/mL penicillin, 100 µg/mL streptomycin and 400 µM 1-thioglycerol supplemented with 50 µg/mL ascorbic acid, 25 ng/mL recombinant human BMP4 and 3 µM CHIR99021 together with 10 µM Rock Inhibitor Y-27632 for the first 24 hours[32]. For cell-cycle and proliferation analysis *via* flow cytometry, iPSCs and CPs were dissociated using Accutase. For autophagy stimulation, CPs were starved or treated at day 3 with 10 µg/ml brefeldin-A for 6 hours. Inhibition of autophagy in control CPs was carried out by incubation at day 1.5 with 100 µM chloroquine (Sigma-Aldrich) for 12 hours. For EIF2AK3 (PERK) activation, CPs were incubated for 6 hours with 5 µM EIF2AK3 activator (CCT020312) at day 1.5 or day 3, as indicated.

For MS-based proteomic analysis and RNAseq, iPSCs were differentiated according the protocol of[54]. iPSCs were dissociated using 0.5 mM EDTA/PBS solution and seeded as single cells on matrigel-coated plates in TeSR™-E8™ until confluency attained 85-95%. On day 0, medium was exchanged to CDM3 (RPMI 1640 supplemented with 500 µM/mL *Oryza sativa*-derived recombinant human albumin and 213 µg/mL L-ascorbic acid 2-phosphate) supplemented with 4-6 µM CHIR99021 (Selleckchem). On day 2, CDM3 medium was supplemented with 2 µM Wnt-C59. On day 4, medium was exchanged to CDM3 medium without supplements, being replaced every other day[54].

For down-stream experiments, beating regions were manually dissected on day 15-18 and reseeded on fibronectin-coated dishes in EB2 medium (DMEM/F12 containing 2% FBS, 1x MEM nonessential amino acids, 1x penicillin/streptomycin, 0.1 mM β-mercaptoethanol and 2 mM L-glutamine). CMs were dissociated at day 60 into single cells using 480 U/mL collagenase type II (Worthington) for assessment of cell-cycle phases and apoptosis *via* flow cytometry. For immunocytochemical analysis of CM-subtypes, multinucleation, and proliferation, CMs were dissociated at day 27 and day 57, plated on fibronectin-coated slides in EB2 medium, and fixed on day 30 and 60, respectively. For analysis of sarcomere length and CM classification on day 60, CMs were dissociated using 10x TripLE Select (Thermo Fisher Scientific) for 15 min at 37°C and plated on micropatterned slides (4Dcell) at day 55. CMs were treated for 30 min with 15 mM 2,3-butanedione-2-monoxime (BDM) to induce relaxation and directly fixed for staining.

For scRNAseq, iPSCs were dissociated using 0.5 mM EDTA/PBS solution and seeded as single cells on matrigel-coated plates in TeSR™-E8™ until confluency attained 85-95%. On day 0, cells were treated with 4-6 µM CHIR99021 (Selleckchem) in RPMI 1640 supplemented with B27 minus insulin. On day 3, cells were treated with 2 µM Wnt-C59 (Selleckchem) in RPMI 1640 supplemented with B27 minus insulin. By day 7, beating CMs were observed and cells were switched to RPMI supplemented with B27. Cultures were subjected to 2-4 days of glucose deprivation starting on day 8 or 9 of differentiation to enrich for cardiomyocytes.

**iPSC-CM-derived 3D heart tissue generation and culture**

iPSC-CM-derived 3D cardiac tissues were generated by seeding iPSC-CMs on decellularized slices of non-human primate LV myocardium. Non-human primate cardiac tissue samples were shipped from the “Deutsches Primatenzentrum Göttingen” where the primates had been euthanized in the course of a vaccination study (file reference: 33.19-42502-04-16/2264). Myocardial tissue was obtained from left mid-ventricular transmural sections and immediately placed in a 30 mM BDM solution at 4°C. The sections were embedded in 4% agarose and further processed to 1.0 × 0.5 cm myocardial tissue slices with 300 µm thicknesses by vibratome cutting (VT1200S, Leica Biosystems, Wetzlar, Germany). Freshly cut myocardial tissue slices were washed twice with PBS and incubated in lysis buffer (10 mM Tris suspended in 0.1% EDTA, pH 7.4) at room temperature for 2 hours. Sections were then incubated in solubilization solution (0.5% sodium-dodecylsulfate (SDS) in PBS) for 6 hours with orbital shaking at room temperature. After this step, samples were washed three times with PBS (containing 0.9 mM calcium and 0.5 mM magnesium; $PBS^{+/+}$) for 10 min, incubated in $PBS^{+/+}$ overnight, and then incubated in FBS for another 12 hours at 37°C. For re-cellularization, cardiac ECM (cECM) scaffolds were anchored in biomimetic culture chambers *via* small plastic triangles attached to the slices with histoacryl according to the fiber direction. iPSC-derived CMs were dissociated at day 15 into single cells using 480 U/mL collagenase type II. After suspension in 0.4% collagen solution (Biochrome), $2 \times 10^6$ cells/slice were seeded using a bioprinting device equipped with a 0.58 mm Luer-lock nozzle (Vieweg Dosier- und Mischtechnik, Kranzberg, Germany). Newly generated 3D heart patches were kept in EB20 medium (same composition as EB2 medium but containing 20% FBS) for 2 days and then in EB2. One week after re-cellularization, 3D cECM heart patches were transferred to custom-made biomimetic culture chambers[45], subjected to a physiological preload of 1 mN and electrical stimulation at 1 Hz (50 mA pulse current, 1 ms pulse duration), and maintained at 37°C, 5% $CO_2$, 20% $O_2$, and 80% humidity. Contraction force and beating rate of the constructs were constantly measured up to 24 days. Contractility data were imported into and analyzed by LabChart Reader software.

**Whole-exome sequencing (WES)**

Exome-enriched indexed libraries were constructed with the SureSelect All Exon system (Agilent Technologies) following manufacturer's recommendations. Paired-end (100 base pair [bp]) sequencing was performed on HiSeq2500 or Hiseq4000 platform (Illumina, San Diego CA). The raw data were processed with an internal bioinformatics pipeline with established methods[55]. In brief, read-mapping to the reference genome (hg19) was done with the Burrows-Wheeler Aligner (BWA) tool. Single-nucleotide variants and small insertions and deletions were called with SAMtools and PINDEL, and all synonymous and intronic (other than canonical splice sites) variants were excluded. Functional annotation was performed using custom scripts on the UCSC gene definition set for hg19. Variants satisfying the criteria of a Phred-scaled genotype quality ≥30, mapping quality ≥50, and a read depth ≥10 were further analyzed to assess their *de-novo* status. Genetic variants were interrogated in the 1000 Genomes project (www.1000genomes.org), the Exome Aggregation Consortium (ExAC) browsers (http://exac.broadinstitute.org), and Genome Aggregation Database (https://gnomad.broadinstitute.org/); a MAF filter was set to be < 0.0001. Predicted functional effect of a coding variant was surveyed using Polyphen2 (http://genetics.bwh.harvard.edu/pph/), SIFT (http://sift.jcvi.org/), and Combined Annotation Dependent Depletion (CADD; http://cadd.gs.washington.edu/). The raw read data of the variants identified was then manually investigated using the Integrative Genomics Viewer (IGV). At a later stage, following technological advancements all the samples were reanalyzed using BWA-Mem algorithm and the Genome Analysis ToolKit GATK software (https://gatk.broadinstitute.org) according to the established GATK Best Practices (https://gatk.broadinstitute.org/hc/en-us/sections/360007226651-Best-Practices-Workflows), but among the additional *de-novo* mutations discovered none passed quality control. Notably also no variant analyzed with PINDEL resulted in a TP *de-novo*. Sanger sequencing was used to confirm the identified *de-novo* mutations and test the carrier status of unaffected parents.

**Cardiomyocyte nuclei isolation and nuclear RNA sequencing (nRNAseq)**

Nuclei were isolated from heart tissues as described previously[56]. RNAsin (80 U/mL, Promega) was added to all buffers. Isolated nuclei were stained in 500 µL staining buffer (PBS containing 1% bovine serum albumin (BSA), 22.5 mg/mL glycine, 0.1% Tween-20) using anti-PLN antibody (1:500, A010–14, Badrilla) for 30 min. Subsequently, the corresponding AlexaFluor488-labeled secondary antibody (1:1000, A11029, Thermo Fisher Scientific) was added. After 30 min of incubation, nuclei were pelleted by centrifugation (1000 ×*g*, 5 min) and resuspended in 1 mL PBS containing 1 mM EDTA. Nuclei were filtered (CellTrics 30 µm, Sysmex) and incubated with Draq7 (final concentration 2.25 nM, Cell Signaling) for 10 min. Nuclei were analyzed and sorted by flow cytometry (Bio-Rad S3, Bio-Rad). All steps were performed at 4°C to ensure integrity of chromatin and RNA.

Nuclear RNA was extracted from sorted nuclei using either the RNeasy Micro Kit or the AllPrep DNA/RNA Micro Kit (Qiagen) including on-column DNAse digestion (Qiagen). RNA was reverse transcribed and amplified with the Nugen Ovation RNAseq System V2 (Nugen). The resulting cDNA was fragmented (Bioruptor, Diagenode). Sequencing libraries were constructed from 100 ng fragmented cDNA using the NEBNext Ultra DNA Library Prep Kit (NEB). The necessary PCR cycles were determined by qPCR after addition of Eva Green (Biotium). All libraries were sequenced on Illumina sequencers in paired-end mode. Quality and adapter trimming of sequencing reads was performed prior to mapping to remove low quality reads and adapter contaminations. RNAseq data were mapped to the human genome using STAR v2.4.2a. PCR duplicates were removed using *SAMtools* (http://samtools.sourceforge.net.). For data normalization and differential gene expression *HTSeq count* (default parameters) and *DESeq2* v1.20.0 (default parameters) with a 5% false-discovery rate (FDR) were used.

**RNA sequencing (RNAseq)**

Transcriptome profiling was performed by RNAseq on pooled RNA isolated from iPSCs, iPSC-derived CPs (at day 1, 2, and 3) and iPSC-derived CMs (at day 6, 8, and 14) obtained from three independent differentiation experiments of three different clones per line carried out in triplicates. Total RNA was isolated using the Absolutely RNA Microprep Kit (Agilent

Technologies) and RNA quality was assessed on Agilent 2100 Bioanalyzer (Agilent Technologies) using Agilent RNA 6000 Nano Kit (Agilent Technologies). RNA-sequencing libraries were prepared from 1 µg RNA using TruSeq Stranded Total RNA Library Prep (Illumina). Samples were sequenced on the Illumina HiScan-SQ (Illumina).

The reads were filtered for low-quality, contaminating 5′ adapters, homopolymers and trimmed for 3′ adapters. Quality control analysis was performed using *FastQC*, trimming of small 3' RNA adapter sequences, using *Trimmomatic v0.36* with default parameters. Quality checked reads were then aligned to the human genome (GRCh37 assembly) using *TopHat2* (default parameters). Gene annotation was obtained for all known genes in the human genome, as provided by Ensemble (GRCh37) (https://support.illumina.com/sequencing/sequencing_software/igenome.ilmn). Using the reads mapped to the genome, the number of reads mapping to each transcript was calculated with *HTSeq-count* (default parameters). Raw read counts were then used as input to *DESeq2* v1.20.0. (default parameters) for calculation of size factor and scaling factor of normalized signal to bring the count values across all the samples to a common scale for each transcript. For the analysis, genes with an adjusted *p*-value < 0.05 were deemed DEGs.

**WES, nRNAseq, and RNAseq data analyses**

For burden testing of GO pathways enriched by D-DNMs, *de-novo* counts from both HLHS cohorts (Munich and PCGC) were merged. This choice was made based on the relatively small size of the Munich cohort and confidently supported by both the similar analysis approach (following GATK Best Practices[27]) and the comparable NS-DNM ratios and distributions of variant classes between the two cohorts (Supplemental Figure 1B).

Evaluation was conducted using R, and a one-tailed Fisher's exact test was performed. Odds-ratios (OR) and their relative *p*-values were calculated[57]. Where ORs could not be calculated we applied the Haldane correction, by setting the null counts of the Fisher's test to 0.5. Enrichment test on the pathways was performed for LOF and D-Mis variants. For randomization test, $10^5$ cycles of randomization of the dataset were run, by shuffling the variants and keeping the number of cases and controls constant, and OR distribution and its standard deviation were calculated. The *p*-values reported represent the probability of the real data's ORs under the null hypothesis (of no enrichment).

Functional interaction networks of D-DNM genes were performed using the Netbox software[58]. Connecting input genes in a network, Netbox partitions the network in modules and identifies significant linker genes. Netbox reports modularity score and an empirical P value of the network build.

Functional interaction networks of dysregulated genes involved in cell-cycle were analyzed using NetworkAnalyst 3.0 (https://www.networkanalyst.ca/NetworkAnalyst/faces/home.xhtml) and the STRING Interactome with a confidence score cutoff of 900.

Fuzzy clustering of DEGs among fetal, infant and adult sample was performed using Mfuzz R package[59], based on Euclidean distance and the c-means. For clustering, gene function and annotation to GO terms were considered.

Gene Set Enrichment Analysis (GSEA) was performed using an algorithm implemented in the R Software Environment for Statistical Computing. Statistical significance was assessed through data set randomization by permuting gene sets 1000 times and considering only gene sets with adjusted *p*-value <0.05.

Enrichment map of gene ontology (GO) terms from differentially expressed genes (DEGs) was performed with *Cytoscape* 3.6.1 using *BINGO* 3.3.0 database[60].

**Single-cell RNA sequencing (scRNAseq) and data analyses**

For scRNASeq, CMs were dissociated at day 14 of differentiation and dispersed into single cell suspension using 10x TrypLE Select (Thermo Fisher Scientific), filtered through a 70 µm filter and resuspended in RPMI 1640 medium supplemented with B27, and submitted for single cell RNA sequencing. After counting, 5,000 cells for each sample were processed using Chromium Single Cell 3’ Library & Gel Bead Kit v2 (10x Genomics) Chromium Single Cell A Chip Kit (10x Genomics) and Chromium i7 Multiplex Kit (10x Genomics) to generate Gel Bead-In-EMulsions (GEMs) and single cell sequencing libraries. Libraries were pooled and sequenced using the HiSeq 4000 (Illumina, San Diego, CA) with a read depth of at least 25,000 reads per cell.

Raw RNA UMI counts were normalized, centered, and scaled before regression was used to remove undesired sources of variability using the standard Seurat R package workflow. These pre-processed data were then analyzed to identify variable genes[61], which were used to perform canonical correlation analysis (MultiCCA). Statistically significant PCs were selected by PC elbow plots and used for t-SNE analysis. Clustering parameter resolution was set to 0.6 for the function FindClusters in Seurat. DEGs were obtained using Wilcoxon rank sum test using as threshold $p$-value $\leq$ 0.05. We used adjusted p-value based on Bonferroni correction using all features in the dataset. For the cell type-specific analysis, single cells of each cell type were identified using FindConservedMarkers function as described within Seurat pipeline. GO analysis of DEGs was performed with ToppGene[62].

Pseudotime Analysis was performed with Monocle2 package[63]. The pseudotime was calculated by selecting the DEGs (q <0.05) to construct the minimal spanning trees (MST). DEGs across the pseudotime axis were identified using a prebuild function in Monocle packages. Second level of pseudotime analysis was performed to catch the influences of other cells. Cells were selected from the corresponding subclusters described in[64] and after normalization and batch correction a trajectory analysis was performed using the standard parameters.

For cell-cycle analysis, a previously defined core set of 43 G1/S and 54 G2/M cell-cycle genes was used[65]. We defined the approximate cell-cycle state of each cell with the expression of the genes in these two gene sets using a function implemented in *yaGST* R package (https://rdrr.io/github/miccec/yaGST/). The single cells highly expressing G1/S or G2/M phase markers were scored as proliferative and the single cells not expressing any of these two categories of genes were scored as not proliferative. Similar approach was used for the other gene signatures analyzed. For the “CM Maturation Signature” and the “Human Fetal Heart Signature” gene sets from DeLaughter et al.[66] and from Sahara et al.[28] were used, respectively. For the “Human Fetal LV/RV signatures” we considered LV and RV enriched genes with expression fold-chance >1.5 from Cui et al.[29], whose ventricular chamber specific expression could be validated manually using the 3D viewer of the human embryonic heart from Asp et al.[43] (LV: *CXCL14*, *HAND1*, *SLIT2*, *IRX3*, *GJA1*, *CRNDE*; RV: *RXRG*, *GPR22*, *GPRIN3*, *C7*).

**MS-based proteomic analysis**

Proteome profiling was performed on isolated iPSCs-derived CMs at day 8 and 14 of cardiac differentiation. Samples were prepared following the in-StageTip sample preparation protocol with minor modifications[67]. Briefly, cells were resuspended in the reducing alkylating sodium deoxycholate buffer (PreOmics) before protein denaturation at 100°C for 10 min. Samples were further sonicated 15 min using a Biorupter (30 s on/off cycles, high settings). Samples were then incubated with LysC and trypsin (1:100, Wako, Sigma-Aldrich) for overnight digestion at 37°C. Resulting peptides were acidified to a final concentration of 0.1% trifluoroacetic acid (TFA) for SDB-RPS binding and desalted before LC-MS/MS analysis.

MS analyses were performed on a quadrupole Orbitrap mass spectrometer[68,69] Q Exactive HF, Thermo Fisher Scientific) coupled to an EASYnLC 1200 ultra-high-pressure system (Thermo Fisher Scientific) via a nano-electrospray ion source. About 0.5 µg of peptides were loaded on a 40 cm HPLC-column (75 µm inner diameter; in-house packed using ReproSil-Pur C18-AQ 1.9 µm silica beads; Dr Maisch GmbH, Germany).

Peptides were separated using a linear gradient from 3% to 23% B in 82 min and stepped up to 40% in 8 min at 350 nL per min where solvent A was 0.1% formic acid in water and solvent B was 80% acetonitrile and 0.1% formic acid in water. The total duration of the gradient was 100 min. Column temperature was kept at 60°C. The MS was operated in 'top-15' data-dependent mode, collecting MS1 spectra with 60,000 resolution, 300-1,650 m/z range and an automatic gain control (AGC) target of 3E6 and a maximum ion injection time of 25 ms. The most intense ions from the full scan were isolated with a width of 1.4 m/z and higher-energy collisional dissociation (HCD) was set with a normalized collision energy (NCE) of 27%. MS2 spectra were collected in the Orbitrap (15.000 resolution) with an AGC target of 1E5 and a maximum ion injection time of 25 ms. Precursor dynamic exclusion was enabled with a duration of 30 s.

MS data were blasted against the 2015 Uniprot human databases (UP000005640_9606 and UP000005640_9606_additional) in the MaxQuant environment using version 1.5.3.34 with a 1% FDR at the peptide and protein level. Peptides with a minimum length of seven amino acids with carbamidomethylation as a fixed modification and protein N-terminal acetylation and methionine oxidations as variable modifications[70]. Enzyme specificity was set as C-terminal to arginine and lysine using trypsin as protease and a maximum of two missed cleavages were allowed in the database search. The maximum initial mass tolerance for precursor and fragment ions were 4.5 ppm and 20 ppm, respectively. Matching between runs was activated with a 0.7 min window. Label-free quantification was performed using a minimum ratio count of 1.

Mass spectrometry proteomics data were analyzed with DEP package (https://rdrr.io/bioc/DEP/) using the functions provided for data normalization, preparation, filtering, variance normalization and imputation of missing values, as well as statistical testing of differentially expressed proteins. Functional interaction networks of dysregulated proteins were performed using NetworkAnalyst 3.0 (https://www.networkanalyst.ca/) and the STRING Interactome with a confidence score cutoff of 900.

**Karyotype analysis**

Karyotype analysis was performed at the Institute of Human Genetics of the Technical University of Munich using standard methodology. Karyotype-specific anomalies, known to occur in human iPSCs[71,72] were tested with the hPSC Genetic Analysis Kit (StemCell$^{TM}$ Technologies) according to the manufacturer´s instructions. Evaluation was performed using the Genetic Analysis App (https://stemcell.shinyapps.io/psc_genetic_analysis_app/).

**Cell-cycle, apoptosis, proliferation, and autophagy analysis by flow cytometry**

Cell-cycle analysis was based on the quantification of DNA content by propidium iodide (PI) using flow cytometry. iPSCs, CPs (at day 1, 2, and 3) and CMs at day 60 were dissociated and fixed with ice-cold 70% ethanol for two hours on ice. Cells were then washed three times with PBS and incubated for 30 minutes with 200 μg/mL RNAse A (Qiagen) at room temperature. Subsequently, 1 mg/mL PI was added for 30 minutes at room temperature and samples were analyzed by flow cytometer. Prior to RNAse A treatment, CMs were washed three times with PBS and stained with rabbit anti-cTNT (Abcam) for one hour at room temperature, followed by incubation with secondary anti-rabbit antibody conjugated with AlexaFluor647. To exclude cell doublets, single cells were identified with forward scatter versus side scatter and subsequent pulse shape analysis applying pulse area versus pulse width on the gated single cells. The G1 phase is characterized by the lower PI signal due to the lower DNA content, while the G2/M phase is characterized by double DNA content corresponding to a higher PI signal intensity.

For quantification of proliferation, iPSCs and CPs (at day 1, 2, and 3) were incubated with 10 µM EdU for two hours, dissociated, fixed with 4% paraformaldehyde (PFA) for 15 min at room temperature, washed three times with PBS and processed using the Click-iT EdU Alexa Fluor 488 Flow Cytometry Assay Kit (Thermo Fisher Scientific) according to manufacturer's instructions. The samples were then subjected to immunostaining with rabbit anti-cleaved caspase 3 antibody (Cell Signaling Technology) overnight at 4°C, followed by incubation with secondary antibody anti-rabbit AlexaFluor-Pacific-Blue for one hour at room temperature.

For assessment of apoptosis in CMs at day 60, cells were dissociated, fixed with 4% PFA for 15 min, and co-immunostained with rabbit anti-cleaved caspase 3 and mouse anti-cTNT antibodies (Thermo Fisher Scientific) overnight at 4°C, followed by incubation with secondary antibodies anti-rabbit AlexaFluor-Pacific-Blue and anti-mouse AlexaFluor647 for one hour at room temperature.

For detection of autophagic flux, CPs were dissociated at day 3 and processed using Cyto-ID™ Autophagy Detection Kit 2.0 (Enzo Life Sciences) according to manufacturer's instructions. The autophagy activity factor (AAF) was calculated based on mean fluorescence intensity of autophagic vacuoles as described[73].

Flow cytometry acquisition analysis was performed with Gallios (Beckman Coulter) and data were evaluated with Kaluza software (Beckman Coulter).

**Immunocytochemistry**

iPSCs were fixed with ice-cold methanol (for TRA1-81 and NANOG staining)/acetone (for SOX2 staining) for 10 min at -20°C, washed and permeabilized with PBS/0.1% Triton-X-100 for 10 min. After blocking with 5% goat serum for 30 min at room temperature, samples were subjected to immunostaining using the following antibodies: mouse anti-TRA1-81 (Merck Millipore), rabbit anti-NANOG (Abcam) and rabbit anti-SOX2 (Abcam) in PBS/0.1% Triton-X-100 containing 1.5% goat serum for one hour at room temperature. After washing with PBS/0.1% Triton-X-100, AlexaFluor®488- and AlexaFluor®555-conjugated secondary antibodies (Abcam) specific to the appropriate species were incubated in PBS/0.1% Triton-X-100 containing 1.5% goat serum for one hour at room temperature. Images were taken with an Axiovert 200M (Zeiss) using Carl ZeissTM Axio Vision Rel. 4.8.2 software (Zeiss).

CMs at day 60 were directly fixed with 4% PFA for 15 min at room temperature after relaxation with 2,3-Butanedione 2-Monoxime. CMs were permeabilized with PBS/0.25% Triton-X-100 at room temperature for 10 min. After blocking with PBS/0.1% Triton-X-100 containing 5% goat serum for 60 min at room temperature, samples were stained with mouse sarcomeric α-actinin (Abcam) in PBS/0.1% Triton-X-100 with 1.5% normal goat serum overnight at 4°C. The AlexaFluor555-conjugated secondary antibody (Abcam) was used in PBS/0.1% Triton-X-100 containing 1.5% goat serum for one hour at room temperature. Classification of CMs from controls and HLHS were analysed based on the classification of Ang et al., 2016. Images were taken using an Axiovert 200M microscope (Zeiss) generating Z-stacks.

3D heart patches were fixed with 4% PFA for 2 hours at 4°C, frozen in OCT (Sakura), and sectioned at 20 μm. CPs and CMs were fixed with 4% PFA for 15 minutes at room temperature. The samples were permeabilized and blocked with 10% FBS (CPs) or 10% goat serum (CMs) in PBS/0.1% Triton-X-100 for one hour at 37°C. CPs were stained with primary antibodies for ISL1 (Developmental Studies Hybridoma Bank), NKX2-5 (Novus Biologicals), TBX5 (Abcam) in PBS/0.1% Triton-X-100 containing 1% FBS; CMs and sections were stained with MLC2a (Synaptic Systems), MLC2v (Proteintech), PKP2 (OriGene), cTNT (Abcam and Thermo Fisher Scientific), Ki67 (BD Biosciences), cleaved caspase-3 (Cell Signaling Technology), cleaved caspase-3 (Alexa Fluor® 488-conjugated, R&D Systems), phosphohistone 3 (Cell Signaling Technology), ph-P53 (Abcam), and α-actinin (Sigma-Aldrich) in PBS/0.1% Triton-X-100 containing 1% goat serum overnight at 4°C. AlexaFluor488 - , AlexaFluor594 - , AlexaFluor647-conjugated secondary antibodies (Thermo Fisher Scientific/Jackson Immunoresearch) specific to the appropriate species were used in PBS/0.1% Triton-X-100 containing 1% FBS (CPs) or 1% goat serum (CMs) for one hour at 37°C. Phalloidin-AlexaFluor594 (Thermo Fisher Scientific) was used to stain F-actin. Apoptosis of CMs in section of heart patches was examined by TUNEL assay using the *In situ* Cell Death detection Kit TMR Red (Roche) according to manufacturer's instructions. Viability of 3D heart patches was determined using LIVE/DEAD™ Viability/Cytotoxicity Kit (Thermo Fisher Scientific) according to the manufacturer's instructions. Nuclei were detected with 1 μg/mL Hoechst 33258 (Sigma-Aldrich). Images were acquired using DMI6000-AF6000 and SP8 confocal laser-scanning Leica microscopes (Wetzlar, Germany).

For LC3B staining, CPs were incubated for 6 hours with 50μM chloroquine before fixation with 4% PFA for 10 minutes at room temperature, permeabilized with 100% ice cold methanol for 10 min at −20°C and washed three times with PBS. After blocking with 3% BSA/PBS for 30 minutes at room temperature, samples were incubated with LC3 antibody (1:100) in 3% BSA/PBS over-night at 4°C. Next day, secondary antibody AlexaFluor-488-donkey anti-rabbit IgG (Thermo Fisher Scientific) was incubated in 3%BSA/PBS for 1 hour at room temperature.

Images were acquired using an SP8 II confocal laser-scanning Leica microscope. Images were assigned with pseudo-colors and processed with ImageJ or Adobe Photoshop. For quantification of TBX5, ISL1, NKX2-5 immunolabeled CPs, marker intensity was measured using Image-J software, applying the same acquisition settings and exposure. The cutoffs for high and low were determined according to the mean intensity level of each marker, excluding statistically significant outliers. Area of the LC3-positive puncta was semiautomatically quantified using a NIH ImageJ macro (GFP-LC3) developed specifically for this purpose[74,75]. For each cell, the overall area of LC3-positive puncta was expressed as a percentage of the area of the whole cell.

**Western Blotting**

Protein samples of CPs were prepared using RIPA buffer (Sigma-Aldrich), separated by SDS-PAGE and blotted according to standard protocols. For LC3 detection, cells were incubated for 6 hours with 50μM chloroquine before lysis and a 4-20% polyacrylamide gradient gel (Bio-Rad) was used. Incubation with primary antibodies eIF2α (Cell Signaling Technology), Phospho-eIF2α (Ser51) (Cell Signaling Technology), LC3 (Novus Biologicals), p62 (Enzo Life Sciences), and GAPDH (Cell Signaling Technology) was performed overnight at 4°C. HRP-conjugated secondary antibodies (Jackson Immunoresearch) were applied next day for one hour. Protein band intensity was quantified using ImageJ.

**RNA isolation, reverse transcription PCR (RT-PCR), and quantitative real-time PCR (qRT-PCR)**

For semi-quantitative analysis, RNA was isolated using the peqGOLD Total RNA Kit (VWR International) and 100 ng RNA were reversely transcribed into cDNA using M-MLV Reverse Transcriptase Kit (Thermo Fisher Scientific). One µl of cDNA was subjected to subsequent PCR using *Taq* polymerase.

For qRT-PCR analysis, RNA was isolated using the Absolutely RNA Microprep Kit (Agilent Technologies) and 1 μg was used to synthesize cDNA with the High-Capacity cDNA Reverse Transcription kit (Applied Biosystems). Gene expression was quantified by qRT-PCR using 1 μL cDNA and the Power SYBR Green PCR Master Mix (Applied Biosystems). Gene expression levels were normalized to *GAPDH* or *ACTB*.

**Calcium imaging of CMs within 3D artificial cardiac tissues and of single 2D-differentiated CMs**

The artificial myocardial tissue slices were loaded with 3 µM Fluo-4-AM (Thermo Fisher Scientific) in culture medium supplemented with 0.75% Kolliphor EL and 30 mM 2,3-butanedione-2-monoxime (BDM) by incubation at 37°C for 60 min, washed, and incubated for another 30 min at 37°C to allow de-esterification of the dye in Tyrode's solution supplemented with $Ca^{2+}$ (135 mM NaCl, 5.4 mM KCl, 1 mM $MgCl_2$, 10 mM glucose, 1.8 mM $CaCl_2$, and 10 mM HEPES; pH 7.35) containing 30 mM BDM. For imaging of single 2D-differentiated CMs, cells at day 30 of differentiation were dissociated to single cells and seeded onto 3.5-cm glass-bottomed cell culture microdishes (MatTek Corporation) coated with 2 µg $cm^{-2}$ fibronectin (Sigma–Aldrich) at a density of $5 \times 10^3$ cells per $cm^2$. Ten days after seeding, loading with 2 µM Fluo-4-AM for 30 min at 37 °C, de-esterification of the dye for 30 min at 37 °C and imaging were all performed in Tyrode's solution supplemented with $Ca^{2+}$.

Calcium signals from CMs within the 3D constructs were subsequently imaged in the above-mentioned solution using an upright epifluorescence microscope (Zeiss Axio Examiner) equipped with a 40x objective, a GFP filter set, and a Rolera em-c2 EMCCD camera. Calcium imaging on single 2D-differentiated CMs an inverted epifluorescence microscope (DMI6000B; Leica Microsystems) equipped with GFP filter sets, a HCX PL APO 63×/1.4-0.6 oil immersion objective (Leica Microsystems) and a Zyla V sCMOS camera (Andor Technology).

Field stimulation electrodes were connected to a stimulus generator (HSE Stimulator P, Hugo Sachs Elektronik, March-Hugstetten, Germany) providing depolarizing pulses (50 V, 5 ms duration) at different frequencies as indicated. Imaging settings (illumination intensity, camera gain, binning) were adjusted to achieve an optimal signal to noise ratio while avoiding pixel saturation at an imaging rate of 50 Hz (3D constructs) or 100 Hz (2D-differentiated cardiomyocytes). ImageJ was used to quantify fluorescence over single cells and over background regions. Subsequent analysis was performed in RStudio using custom-written scripts as described previously[52]. After subtraction of background fluorescence, the time course of Fluo-4 fluorescence was expressed either in arbitrary units or normalized to the initial value (F/F0). After manual selection of the starting points and the peaks of the calcium transients, the transient duration at 90% decay (TD90), was automatically determined by the script.

#### SUPPLEMENTAL RESULTS

##### Proteomic analysis confirms cell-cycle defects in HLHS CMs

To validate transcriptional defects of HLHS CMs at the protein level, we performed mass spectrometry-based analysis of iPSC-CMs. In diseased cells, we found 83 differentially expressed proteins (DEPs) (35 up- and 48 down-regulated) at D8 and 51 (25 up- and 26 downregulated) at D14 (Table III and Figure VIIIA and VIIID in the Data Supplement). At D8, 17 DEPs were hubs of a functional network involving pathways implicated in cellular metabolism, mitotic cell-cycle checkpoints/arrest, chromatin modification/chromosome organization, and DNA damage response (Figure VIIIB and VIIIC and Table III in the Data Supplement). Similar pathways were likewise recovered, together with apoptosis, in the functional network of D14 DEPs (Figure VIIIE and VIIIF and Table III in the Data Supplement).

##### Single cell RNAseq of iPSC-CMs and sub-population cluster identification.

We assigned identities to each population by cross-referencing the most highly and uniquely expressed genes in each cluster with known cardiac subtype markers from human and mouse single-cell studies (Figure 5B and Figure IX in the data Supplement)[28,29,41-43]. *MYL3* expression distinguished CMs committed to the ventricular lineage, while *CLU*, *ID1*, and *HES4* transcripts defined CP states. CMs expressing genes enriched in LV (*HAND1*, *SLIT2*, and *GJA1*) were located in cluster 0, while markers preferentially expressed in RV CMs (*IGFBP5*, *IRX1*, *IRX2*, *CPNE5, and RXRG*) highlighted cells mainly represented in cluster 1. This cluster also comprised cells expressing *KCNA5,* an early marker of atrial commitment. Cluster 2 contained AVC CMs characterized by high levels of genes important for cardiac epithelial-mesenchymal transition (*TGFB2*, *WNT2, BMP2, HAS2*) and AVC/valve development *(TBX2, GATA4, RSPO3*). Transcriptome typical of OFT CMs (*HAND2*, *TGFB2*, *CTGF*) was found mainly in cluster 5. Clusters 3 and 8 encompassed maturing and more differentiated CMs, as indicated by high expression of sarcomeric (e.g. *MYH6*, *MYL3*, *MYL4*, *TNNC1, MYBPC3*) and energy metabolism related (*ENO1/3*, *COX5A/B*, *COX6C*, *COX7B/C*) transcripts as well as gene signature typical of maturing CM during early mouse development[66]. Cell-cycle analysis revealed that cells in clusters 4 and 6 were actively proliferating cells (mainly CMs of diverse subtypes) enriched for genes involved in the G1/S (e.g. *E2F1*, *PCNA*) and G2/M (e.g. *KIF14*, *PLK1*) phases, respectively. Genes linked to cytokinesis (e.g. *AURKA/B)* were also highly expressed in cluster 6. The remaining two clusters were CPs, including early precursors (cluster 7) with exclusive expression of *CLU* and later myocytic committed progenitors (cluster 9) with specific gene expression signature of ventricular chamber precursors[28] (Figures 5C and Figure IXA in the Data Supplement).

#### SUPPLEMENTAL TABLES

**Supplemental Table V. List of antibodies and assays used in the study**

| | SOURCE | IDENTIFIER |
|---|---|---|
| **Primary antibodies** | | |
| Anti-Cardiac Troponin T, mouse monoclonal Antibody (13-11) | Thermo Fisher Scientific | MA5-12960 |
| Anti-Cardiac Troponin T, rabbit monoclonal antibody [EPR3696] | Abcam | ab92546 |
| Anti-Cleaved Caspase-3 (Asp175) (5A1E), rabbit monoclonal antibody | Cell Signaling Technology | 9664 |
| Anti-Cleaved Caspase-3, Alexa Fluor® 488-conjugated rabbit monoclonal antibody | R&D Systems | IC835G-025 |
| Anti- eIF2α, rabbit monoclonal antibody | Cell Signaling Technology | 5324 |
| Anti-GAPDH (14C10), rabbit monoclonal antibody | Cell Signaling Technology | 2118S |
| Anti-ISL1, mouse monoclonal antibody | Developmental Studies Hybridoma Bank | 39.4D5 |
| Anti-Ki-67, mouse monoclonal antibody | BD Biosciences | 556003 |
| Anti-LC3B, rabbit polyclonal antibody | Novus Biologicals | NB100-2220 |
| Anti-MLC2a, mouse monoclonal antibody | Synaptic Systems | 311 011 |
| Anti-MLC-2V, rabbit polyclonal antibody | Proteintech | 10906-1-AP |
| Anti-Nanog, rabbit polyclonal antibody | Abcam | ab106465 |
| Anti-NKX2.5, goat polyclonal antibody | Novus Biologicals | NBP1-51953 |
| Anti-p53 (phospho S6) antibody [Y179], rabbit monoclonal antibody | Abcam | ab32132 |
| Anti-p62, rabbit polyclonal antibody | Enzo Life Sciences | BML-PW9860-0100 |
| Anti-Phospho-eIF2α (Ser51), rabbit monoclonal antibody | Cell Signaling Technology | 3398 |
| Anti-Phospho-Histone H3 (Ser10), rabbit polyclonal antibody | Merck Millipore | 06-570 |
| Anti-Plakophilin 2 (PKP2), guinea pig polyclonal antibody | OriGene | AP09554SU-N |
| Anti-PLN, mouse monoclonal antibody | Badrilla | A010-14 |
| Anti-Sarcomeric α-Actinin, mouse monoclonal antibody | Abcam | ab9465 |
| Anti-SOX2, rabbit polyclonal antibody | Abcam | ab137385 |
| Anti-TBX5, rabbit polyclonal antibody | Abcam | ab137833 |
| Anti-TRA1-81, mouse monoclonal antibody | Merck Millipore | MAB4381 |
| Anti-α-Actinin, mouse monoclonal antibody | Sigma-Aldrich | A7811 |

| Critical Commercial Assays | | |
|---|---|---|
| Agilent Absolutely RNA Microprep Kit | Agilent Technologies | 400805 |
| Agilent RNA 6000 Nano Kit | Agilent Technologies | 5067-1511 |
| AllPrep DNA/RNA Micro Kit | Qiagen | 80284 |
| Chromium i7 Multiplex Kit | 10x Genomics | PN-120262 |
| Chromium Single Cell 3’ Library & Gel Bead Kit v2 | 10x Genomics | PN-120237 |
| Chromium Single Cell A Chip Kit | 10x Genomics | PN-120236 |
| Click-iT EdU Alexa Fluor 488 Cell Proliferation Kit for Imaging | Thermo Fisher Scientific | C10337 |
| Click-iT EdU Alexa Fluor 488 Flow Cytometry Assay Kit | Thermo Fisher Scientific | C10425 |
| CYTO-ID® Autophagy detection kit 2.0 | Enzo Life Sciences | ENZ-KIT175 |
| CytoTune™-iPS 2.0 Sendai Reprogramming Kit | Thermo Fisher Scientific | A16518 |
| DNeasy Blood & Tissue Kit | Qiagen | 69506 |
| EvaGreen® Dye | Biotium | 31000 |
| Fast Start High Fidelity PCR System | Sigma-Aldrich | 4738292001 |
| High-Capacity cDNA Reverse Transcription Kit | Thermo Fisher Scientific | 4368814 |
| hPSC Genetic Analysis Kit | StemCell™ Technologies | 07550 |
| In Situ Cell Death detection Kit TMR Red | Roche | 12156792910 |
| LIVE/DEAD™ Viability/Cytotoxicity Kit, for mammalian cells | Thermo Fisher Scientific | L3224 |
| M-MLV Reverse Transcriptase Kit | Thermo Fisher Scientific | 28025013 |
| NEBNext® Ultra™ DNA Library Prep Kit for Illumina | NEB | E7370 |
| Ovation RNA-Seq System V2 | Nugen | 7102 |
| pegGOLD Total RNA Kit | VWR International | 7322871 |
| Power SYBR® Green PCR Master Mix | Applied Biosystems | 4367659 |
| RNasin, Ribonuclease Inhibitor | Promega | N2511 |
| RNeasy Micro Kit | Qiagen | 74004 |
| SureSelect All Exon system | Agilent Technologies | |
| TruSeq® Stranded Total RNA Library Prep Human/Mouse/Rat (48 Samples) | Illumina | 20020596 |

**Supplemental Table VI. List of primers used in the study**

| Gene and use | Primer forward | Primer reverse |
|---|---|---|
| *ACTA2* qRT-PCR | *5‘-GTGATCACCATCGGAAATGAA-3‘* | *5‘-TCATGATGCTGTTGTAGGTGGT-3‘* |
| *ACTB* RT-PCR | *5‘-CCAACCGCGAGAAGATGA-3‘* | *5‘-CCAGAGGCGTACAGGGATAG-3‘* |
| *AFP* qRT-PCR | *5‘-GTGCCAAGCTCAGGGTGTAG-3‘* | *5‘-CAGCCTCAAGTTGTTCCTCTG-3‘* |
| *AIM1L* PCR | *5’-CTCCTGCTCCATCCACTACAAG-3’* | *5’-CCTTCACAACCTCTTTCTGGGT-3’* |
| *ATF4* qRT-PCR | *5’-TTCTCCAGCGACAAGGCTAAGG-3’* | *5’-CTCCAACATCCAATCTGTCCCG-3’* |
| *ATF6* qRT-PCR | *5’-CAGACAGTACCAACGCTTATGCC-3’* | *5’-GCAGAACTCCAGGTGCTTGAAG-3’* |
| *ATG12* qRT-PCR | *5’-GGGAAGGACTTACGGATGTCTC-3’* | *5’-AGGAGTGTCTCCCACAGCCTTT-3’* |
| *ATG3* qRT-PCR | *5’-ACTGATGCTGGCGGTGAAGATG-3’* | *5’-GTGCTCAACTGTTAAAGGCTGCC-3’* |
| *ATG5* qRT-PCR | *5’-GCAGATGGACAGTTGCACACAC-3’* | *5’-GAGGTGTTTCCAACATTGGCTCA-3’* |
| *BAI2* PCR | *5’-CAAACTCTTCAAGAGTGAGGT-3’* | *5’-TCAACAACAACAACTCTAGCC-3’* |
| *c-MYC* RT-PCR | *5‘-CACCAGCAGCGACTCTGA-3‘* | *5‘-GATCCAGACTCTGACCTTTTGC-3‘* |
| *CACNA1C* qRT-PCR | *5’-CAATCTCCGAAGAGGGGTTT-3’* | *5’-TCGCTTCAGACATTCCAGGT-3’* |
| *CASQ2* qRT-PCR | *5’-CCGGGACAATACTGACAACC-3’* | *5’-CTTCTCCCAGTAGGCAACGA-3’* |
| *CD31* qRT-PCR | *5‘-ATGCCGTGGAAAGCAGATAC-3‘* | *5‘-CTGTTCTTCTCGGAACATGGA-3‘* |
| *CHOP* qRT-PCR | *5’-GGAACCTGAGGAGAGAGTGTTC-3’* | *5’-TTTTGGAAAAGGGTAGGTTAAGTTT-3’* |
| *DENND5B* PCR | *5’-ACATTCTGTGCTGCTAGAGCAT-3’* | *5’-TGATCAGCATGACCGTTCTAG-3’* |
| *EIF2AK3 (PERK)* qRT-PCR | *5’-GTCCCAAGGCTTTGGAATCTGTC-3’* | *5’-CCTACCAAGACAGGAGTTCTGG-3’* |
| *FABP5* qRT-PCR | *5’-CCCTGGGAGAGAAGTTTGAAG-3’* | *5’-ATCCCACTCCTGATGCTGAA-3’* |
| *FGFR1* PCR | *5’-CTATGCTTGCGTAACCAGCAG-3’* | *5’-TTCTCCACAATGCAGGTGTAG-3’* |
| *GAPDH* qRT-PCR | *5‘-TCCTCTGACTTCAACAGCGA-3‘* | *5‘-GGGTCTTACTCCTTGGAGGC-3‘* |
| *GJA1* qRT-PCR | *5’-CACTTGGCGTGACTTCACTACTT-3’* | *5’-CCAGCAGTTGAGTAGGCTTGA-3’* |
| *GJA5* qRT-PCR | *5’-AATCTTCCTGACCACCCTGCATGT-3’* | *5’-CAGCCACAGCCAGCATAAAGACAA-3’* |
| *HEY2* qRT-PCR | *5’-AAGATGCTTCAGGCAACAGG-3’* | *5’-TCATGAAGTCCATGGCAAGA-3’* |
| *ISL1* qRT-PCR | *5‘-AAAGTTACCAGCCACCTTGGA-3‘* | *5‘-ATTAGAGCCCGGTCCTCCTT-3‘* |
| *KCNA5* qRT-PCR | *5’-AGTGTAACGTCAAGGCCAAGAGCA-3’* | *5’-TCACAAATCTGTTTCCCGGCTGGT-3’* |
| *KCNJ3* qRT-PCR | *5’-TCATCAAGATGTCCCAGCCCAAGA-3’* | *5’-CACCCGGAACATAAGCGTGAGTTT-3’* |
| *KLF4* RT-PCR | *5‘-TCTTCGTGCACCCACTTGGG -3‘* | *5‘-CTGCTCAGCACTTCCTCAAG -3‘* |

| | | |
|---|---|---|
| *KMT2A* PCR | *5’-GGAGAGGATGAGCAATTCTTAG-3’* | *5’-CTTGCTGAAACGTAGCAGAAG-3’* |
| *KRT14* qRT-PCR | *5‘-CACCTCTCCTCCTCCCAGTT-3‘* | *5‘-ATGACCTTGGTGCGGATTT-3‘* |
| *MACF1* PCR | *5’-CAATAAGAGTGCCACCGTGGA-3’* | *5’-ATGATGGAAGGATCAATGCCTG-3’* |
| *MYRF* PCR | *5’-CACGATAACCCCTGAGACACTG-3’* | *5’-CTCATGATCTGGACGGGACATT-3’* |
| *NANOG* RT-PCR | *5‘-TGCTTTGAAGCATCCGACTGT-3‘* | *5‘-GGTTGTTTGCCTTTGGGACTG -3‘* |
| *NDFB10* PCR | *5’-CTCGGCGTCCTCTGTAGCG-3’* | *5’-GGCAGGGTGTGGGATCCAG-3’* |
| *NKX2-5* qRT-PCR | *5`-CACCGGCCAAGTGTGCGTCT-3’* | *5‘-GCAGCGCGCACAGCTCTTTC-3’* |
| *OCT4* RT-PCR | *5‘-GGGATGGCGTACTGTGGG-3‘* | *5‘-GCACCAGGGGTGACGGTG-3‘* |
| *PLN* qRT-PCR | *5’-TCCCATAAACTGGGTGACAGA-3’* | *5’-TGATACCAGCAGGACAGGAAG-3’* |
| *REX1* RT-PCR | *5‘-AGTAGTGCTCACAGTCCAGCAG-3‘* | *5‘-TGTGCCCTTCTTGAAGGTTT-3‘* |
| *RYR2* qRT-PCR | *5’-GCTATTCTGCACACGGTCATT-3’* | *5’-ATTTCCGTGCCACTTCCTTT-3’* |
| *SERCA2* qRT-PCR | *5’-ACAATGGCGCTCTCTGTTCT-3’* | *5’-ATCCTCAGCAAGGACTGGTTT-3’* |
| *SeV* RT-PCR | *5‘-GGATCACTAGGTGATATCGAGC-3‘* | *5‘-ACCAGACAAGAGTTTAAGAGATATGTATC-3‘* |
| *SLC8A1* qRT-PCR | *5’-GAGACCTGGCTTCCCACTTT-3’* | *5’-ATTCCCAGGAAGACATTCACC-3’* |
| *SOX2* RT-PCR | *5‘-AGCAGACTTCACATGTCCCAG-3‘* | *5‘-ACCGGGTTTTCTCCATGCTGT-3‘* |
| *SOX7* qRT-PCR | *5‘-TGAACGCCTTCATGGTTTG-3‘* | *5‘-AGCGCCTTCCACGACTTT-3‘* |
| *Spliced-XBP1* qRT-PCR | *5’-TGCTGAGTCCGCAGCAGGTG-3’* | *5’-GCTGGCAGGCTCTGGGGAAG-3’* |
| *SYBU* PCR | *5’-GTCATGAGAGGAGAGTCCACA-3’* | *5’-AGAGGACTGAATGCTTCCTTCA-3’* |
| *TBX5* qRT-PCR | *5‘-GGGCAGTGATGACATGGAG-3‘* | *5‘-GCTGCTGAAAGGACTGTGGT-3‘* |
| *TH* qRT-PCR | *5‘-TGTACTGGTTCACGGTGGAGT-3‘* | *5‘-TCTCAGGCTCCTCAGACAGG-3‘* |
| *Total-XBP1* qRT-PCR | *5’-CTGCCAGAGATCGAAAGAAGGC-3’* | *5’-CTCCTGGTTCTCAACTACAAGGC-3’* |

**SUPPLEMENTAL FIGURES AND FIGURE LEGENDS**

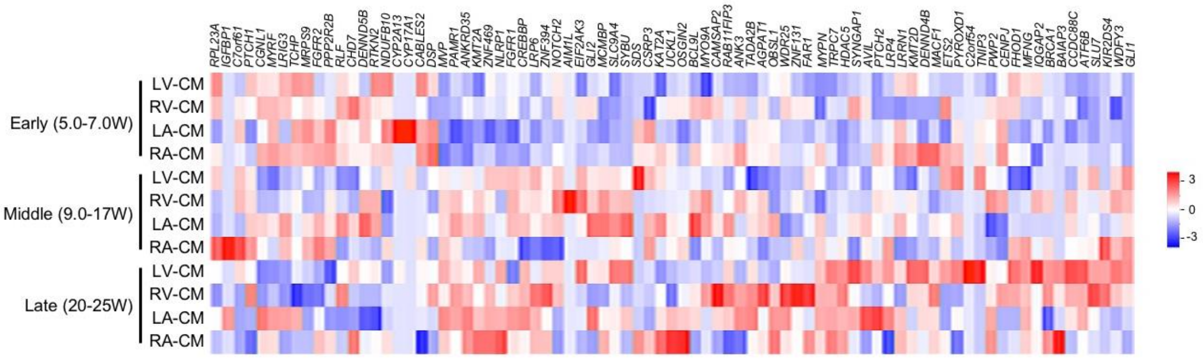


**Supplemental Figure I. Expression of HLHS D-DNM genes in the ventricular and atrial chambers during human heart development (Related to Figure 1).**

Heart chamber-specific expression of HLHS D-DNM genes based on scRNA-seq from normal fetal hearts between 5 and 25 weeks (W) of gestation from Sahara et al[28]. LA, left atrium: RA, right atrium.

A

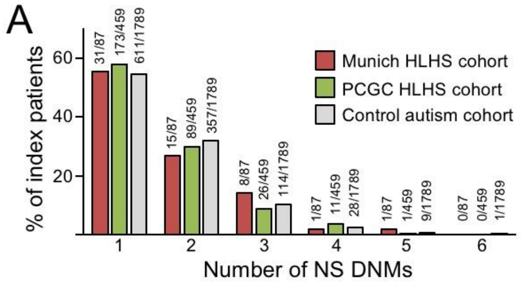


B

| | Munich cohort | PCGC cohort |
|---|---|---|
| HLHS parents-offspring trios | 87 | 459 |
| NS DNMs | 94 | 366 |
| NS DNM ratio | 1.08 | 0.78 |
| Cases with NS DNMs | 56 (64%) | 300 (65%) |
| Cases with LOF DNMs | 12 (14%) | 64 (14%) |
| Cases with Mis DNMs | 50 (57%) | 217 (47%) |

C

| Multi hits in different patients | D-DNM in Munich HLHS cohort | D-DNM in PCGC HLHS cohort |
|---|---|---|
| *CACNA1A* | | p.M1986V<br>p.L1610fs |
| *CHD7* | p.R2027* | p.Q2359* |
| *KMT2D* | p.F3699Lfs*14 | p.V5423F p.R4198*<br>p.R5351Q p.R1757*<br>p.R5154Q p.S1722fs<br>p.N5079fs p.C349*<br>p.F4576fs p.A254fs<br>p.R4536* p.S31* |
| *LRP6* | p.A580Gfs*15<br>p.R360C | |
| *MACF1* | p.A4014L<br>p.A5271P | |
| *MYH6* | | p.Q373P<br>p.R108C |
| *MYRF* | p.I213Tfs*59<br>p.M876Vfs*23 | p.Q394H |
| *RABGAP1L* | | c.-33-1G>A<br>p.S496G |
| *RBFOX2* | | c.1311+1G>C<br>p.R357*<br>p.L299fs |

D

| | Genes (bar; -Log10 p-value, 0–20) | LOF DNM: Hits/Genes in cases | LOF DNM: Hits/Genes in controls | LOF DNM: OR | LOF DNM: P-val | D-Mis DNM: Hits/Genes in cases | D-Mis DNM: Hits/Genes in controls | D-Mis DNM: OR | D-Mis DNM: P-val | LOF + D-Mis DNM (D-DNM): Hits/Genes in cases | LOF + D-Mis DNM (D-DNM): Hits/Genes in controls | LOF + D-Mis DNM (D-DNM): OR | LOF + D-Mis DNM (D-DNM): P-val |
|---|---|---|---|---|---|---|---|---|---|---|---|---|---|
| ***GO Biol Processes*** | | | | | | | | | | | | | |
| Pos regulation of RNA biosynthetic process | 1600 | 28/17 | 15/15 | **4.9** | **5.2e-06** | 34/32 | 19/18 | **3.1** | **1.5e-04** | 62/46 | 34/33 | **3.7** | **1.0e-08** |
| Pos regulation of gene expression | 1974 | 27/16 | 21/20 | **3.2** | **4.3e-04** | 35/33 | 31/29 | **1.9** | **1.6e-02** | 62/46 | 52/48 | **2.3** | **7.3e-05** |
| Pos regulation of transcription by RNA Pol II | 1185 | 23/13 | 11/11 | **5.1** | **2.0e-05** | 27/25 | 10/10 | **4.6** | **2.6e-05** | 50/36 | 21/21 | **4.6** | **4.7e-09** |
| Regulation of cell differentiation | 1863 | 27/15 | 14/14 | **5.0** | **6.0e-06** | 34/30 | 30/29 | **1.9** | **1.7e-02** | 61/44 | 44/43 | **2.7** | **4.4e-06** |
| Circulatory system development | 1161 | 10/9 | 7/7 | **2.9** | **2.9Ee02** | 23/22 | 27/24 | 1.3 | 2.4e-01 | 33/30 | 34/31 | **1.7** | **3.4e-02** |
| Cell projection organization | 1512 | 8/6 | 19/18 | 0.8 | 7.8e-01 | 30/28 | 26/24 | **1.9** | **2.3e-02** | 38/34 | 45/42 | 1.4 | 8.0e-02 |
| Heart development | 573 | 7/6 | 5/5 | 2.8 | 7.3e-02 | 16/15 | 16/14 | 1.5 | 1.7e-01 | 23/20 | 21/19 | **1.9** | **3.5e-02** |
| Cell-cycle | 1847 | 12/12 | 19/18 | 1.2 | 3.7e-01 | 24/24 | 25/23 | 1.5 | 1.3e-01 | 36/35 | 44/41 | 1.4 | 1.1e-01 |
| Cardiac muscle tissue development | 232 | 4/3 | 1/1 | **7.9** | **4.9e-02** | 11/10 | 7/7 | 2.4 | 5.8e-02 | 15/13 | 8/8 | **3.2** | **6.9e-03** |
| Cytoskeleton organization | 1298 | 7/7 | 22/21 | 0.6 | 9.3e-01 | 24/21 | 25/24 | 1.5 | 1.3e-01 | 31/28 | 47/45 | 1.1 | 4.3e-01 |
| Developmental growth | 663 | 15/5 | 8/7 | **4.1** | **1.5e-03** | 22/17 | 11/10 | **3.3** | **1.4e-03** | 37/20 | 19/16 | **3.6** | **8.8e-06** |
| Cell fate commitment | 260 | 3/3 | 2/2 | 2.9 | 2.3e-01 | 10/10 | 3/2 | **5.2** | **7.7e-03** | 13/13 | 5/4 | **4.4** | **3.2e-03** |
| ***Canonical Pathways*** | | | | | | | | | | | | | |
| React_Gli proteins bind promoters of HH responsive genes to promote transcription | 7 | 0/0 | 1/1 | 0.0 | 1.0e+00 | 4/4 | 2/1 | 3.0 | 1.8e-01 | 4/4 | 3/2 | 2.2 | 2.5e-01 |
| KEGG_Notch signaling pathway | 47 | 1/1 | 0/0 | 3.8 | 3.4e-01 | 5/5 | 1/1 | **7.5** | **4.2e-02** | 6/6 | 1/1 | **9.9** | **1.4e-02** |
| React_TCF depend sign in response to Wnt | 232 | 13/4 | 0/0 | **59.1** | **4.9e-07** | 10/8 | 5/5 | **3.1** | **3.4e-02** | 23/10 | 5/5 | **8.1** | **1.4e-06** |
| Reactome_Chromatin organization | 272 | 12/3 | 2/1 | **13.1** | **7.4e-05** | 9/7 | 9/9 | 1.5 | 2.7e-01 | 21/9 | 11/9 | **3.3** | **1.2e-03** |
| Reactome_E2F pathway | 73 | 1/1 | 0/0 | 3.8 | 3.4e-01 | 4/4 | 0/0 | **12.1** | **2.6e-02** | 5/5 | 0/0 | **16.5** | **7.9e-03** |

E

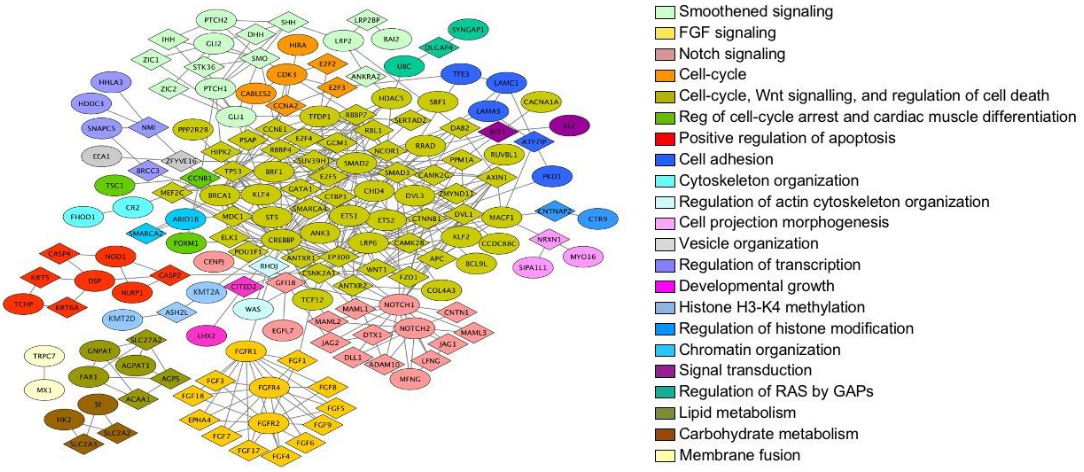


**Supplemental Figure II. Characterization of damaging *de-novo* mutations (D-DNMs) recovered by whole-exome sequencing of the Munich and the PCGC cohorts of HLHS parents-offspring trios (Related to Figure 1). A**, Number of identified *de-novo* nonsynonymous mutations (NS DNMs) per HLHS and control index patients. Controls consist of 1789 trios comprising parents and unaffected siblings of autism probands from[30]. **B**, Distribution of NS DNMs in both HLHS cohorts. LOF and Mis indicate loss-of-function and missense mutations, respectively. **C**, Genes with multiple damaging hits in HLHS cohorts. **D**, Bar chart of Gene Ontology (GO) enrichment analysis of D-DNM genes recovered from both Munich and PCGC HLHS parents-offspring trios (right) and their burden enrichment for the corresponding GO category (left). The number of cases and controls was 546 and 1789, respectively. Odds-ratio (OR) and *p*-value (Fisher's test) are indicated in bold for significant enrichments. OR are Haldane-corrected where appropriate. Numbers in the graph bars indicate total genes in the category. **E**, Protein-protein network analysis of D-DNMs recovered from both HLHS cohorts. Each Netbox module is coded by a different color, with mutated genes illustrated as circle and linker genes as diamonds. Network modularity = 0.51; scaled network modularity = 5.00; random networks modularity = 0.33 ± 0.036 (n = 100, mean ± SD), *p*-value <10e-05.

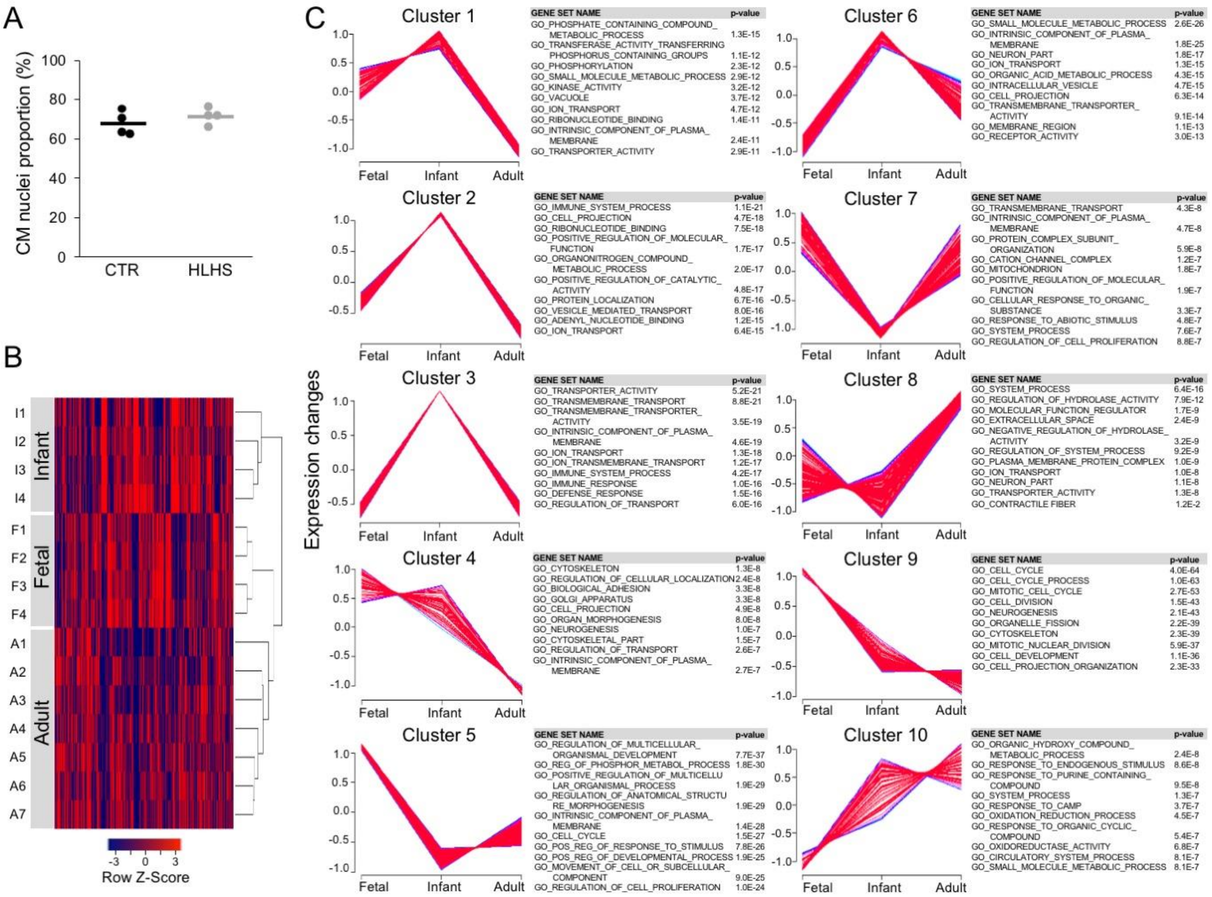


**Supplemental Figure III. Nuclear RNA sequencing of CMs from healthy hearts at 3 developmental stages and from infant HLHS subjects (Related to Figure 2). A**, Percentage of CM nuclei from infant control and HLHS subjects. **B**, Unsupervised hierarchical clustering of the global gene expression data from healthy fetal, infant, and adult CMs. The dendrogram illustrates separation of the samples based on the developmental stage. **C**, Fuzzy clustering showing all DEGs based on their expression at fetal, infant, and adult stages of normal heart development. Top-ten GO terms are shown on the right.

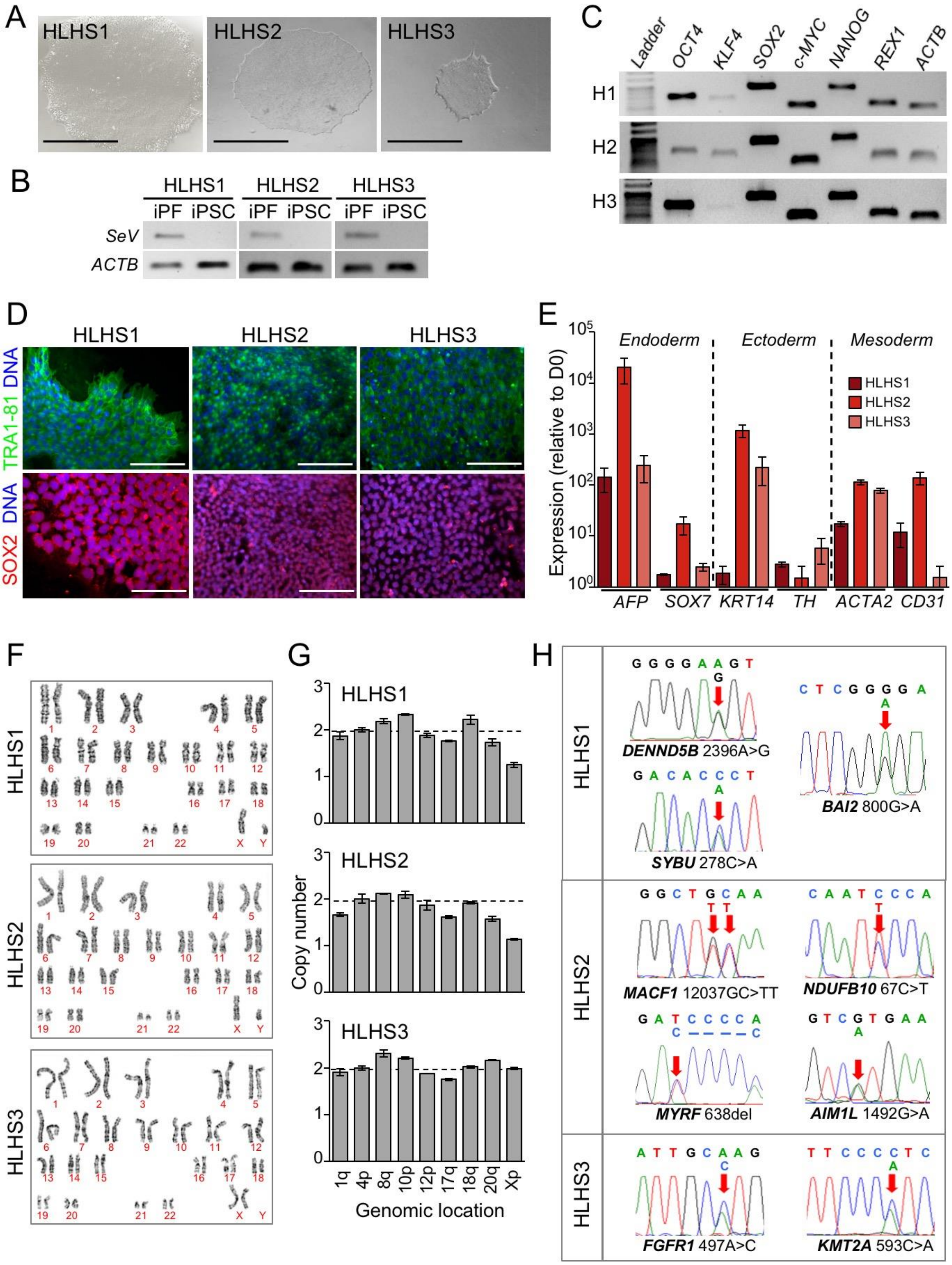


**Supplemental Figure IV. Characterisation of iPSCs from HLHS patients (Related to Figures 3 to 7). A**, Representative iPSC colonies from HLHS1, HLHS2 and HLHS3. Scale bar, 500 µm. **B**, Sendai-footprinting of representative clones from the generated iPSC lines indicating the exclusion of the Sendai virus. iPF, infected parental fibroblasts. **C**, RT-PCR demonstrating the expression of endogenous pluripotency marker genes. **D**, Representative images of iPSC colonies immunostained for TRA1-81 (green) and SOX2 (red). Scale bar, 100 µm. **E**, Quantitative RT-PCR for lineage marker genes representative of the three embryonic germ layers upon spontaneous embryoid-body differentiation (D21 vs D0). Data are mean ± SEM from 3 differentiations per line. **F**, Karyograms of representative clones from the HLHS iPSCs showing normal karyotype. **G**, Quantitative PCR-based genetic analysis of karyotypic abnormalities reported in human iPSCs (hPSC Genetic Analysis Kit, StemCell Technologies) indicating normal copy number over the regions of interest. **H**, Results of Sanger sequencing analysis indicate the presence of the identified heterozygous *de-novo* mutations in the generated iPSCs.

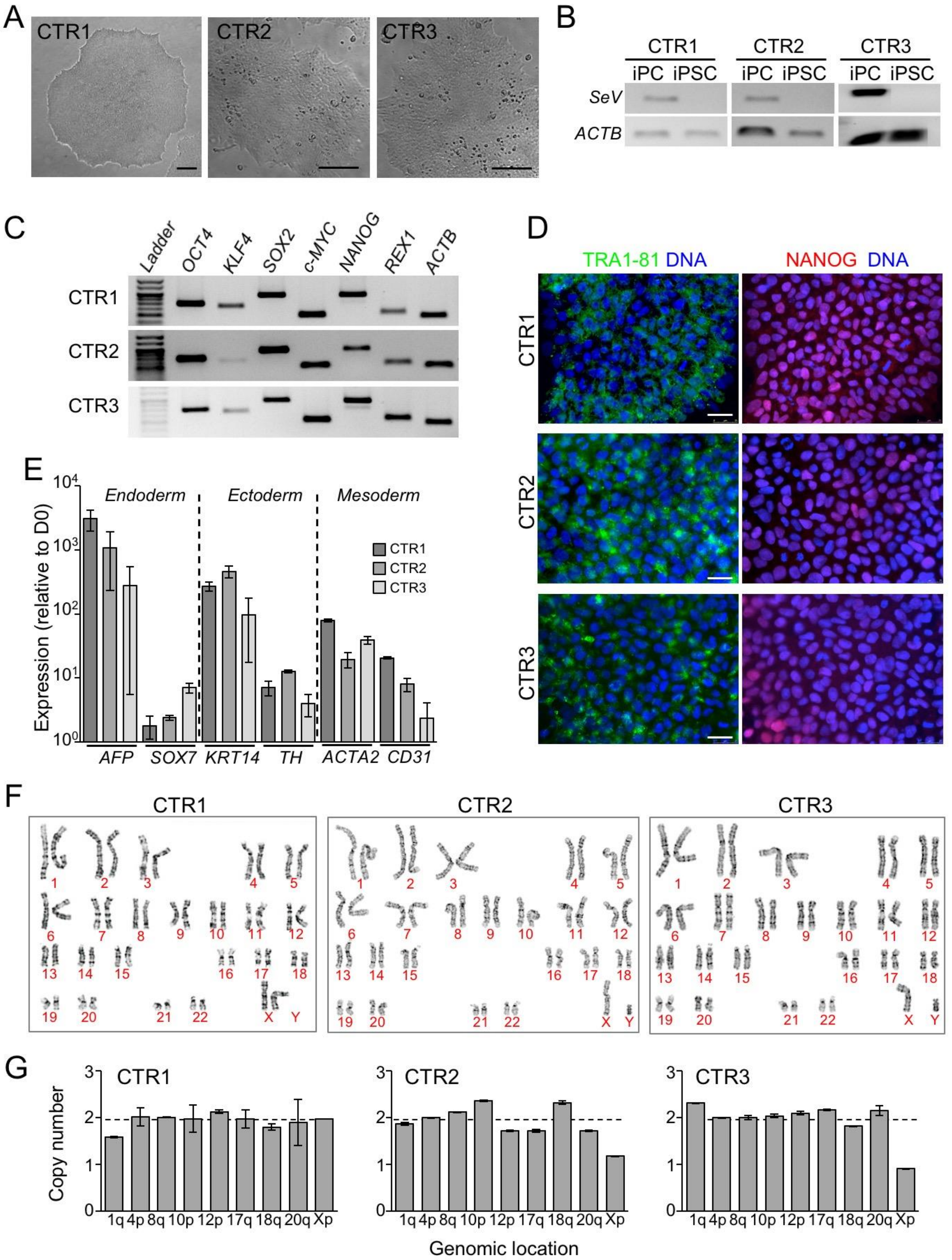


**Supplemental Figure V. Characterisation of iPSCs from healthy control individuals (Related to Figures 3 to 7). A**, Representative iPSC colonies from CTR1, CTR2 and CTR3. Scale bar, 100 µm. **B**, Sendai-footprinting of representative clones from the generated iPSC lines indicating the exclusion of the Sendai virus. iPC, infected parental cells. **C**, RT-PCR demonstrating the expression of endogenous pluripotency marker genes. **D**, Representative images of iPSC colonies immunostained for TRA1-81 (green) and NANOG (red). Scale bar, 25 µm. **E**, Quantitative RT-PCR for lineage marker genes representative of the three embryonic germ layers upon spontaneous embryoid-body differentiation (D21 vs D0). Data are mean ± SEM from 2-3 differentiations per line. **F**, Karyograms of representative clones from the control iPSCs showing normal karyotype. **G**, Quantitative PCR-based genetic analysis of karyotypic abnormalities reported in human iPSCs (hPSC Genetic Analysis Kit, StemCell Technologies) indicating normal copy number over the regions of interest.

A

| KaBi DHM ID / Helmholtz ID | HLHS iPSC | Gene | *De-novo* mutation | Amino acid change | Morphology | | |
|---|---|---|---|---|---|---|---|
| | | | | | LV | AV | MV |
| **375 / 69270** | **HLHS1** | *DENND5B*<br>*SYBU*<br>*BAI2* | missense<br>missense<br>missense | Lys834Arg<br>Pro93Hys<br>Gly267Glu | hypoplastic, thickened | atresia | atresia |
| **612 / 70315** | **HLHS2** | *MACF1*<br>*MYRF*<br>*NDUFB10*<br>*AIM1L* | missense<br>frameshift<br>missense<br>missense | Ala4013Leu #<br>Ile213Thr fs*59<br>Pro23Ser<br>Val498Met | hypoplastic, thickened | atresia | atresia |
| **606 / 73119** | **HLHS3** | *FGFR1*<br>*KMT2A* | missense<br>missense | His166Pro<br>Pro198His | hypoplastic, thickened | atresia | atresia |

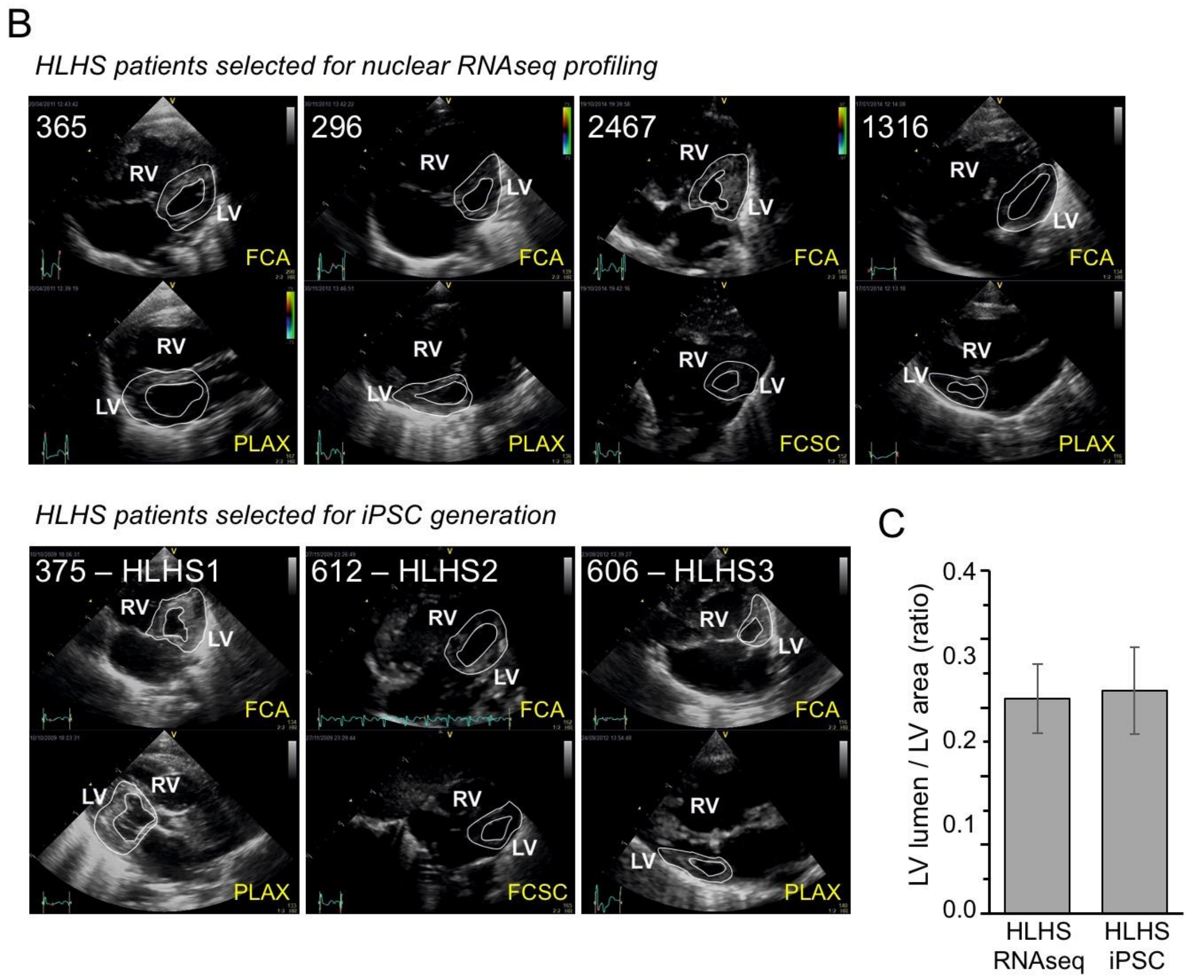


**Supplemental Figure VI. Genetic and echocardiographic information of the HLHS patients selected for iPSC generation (Related to Figures 3 to 7). A**, *De-novo* mutations carried by the three HLHS patients selected for iPSC generation and clinical information. #

two *de-novo* missense mutations in *MACF1* resulted in the amino acid change. AV, aortic valve; LV, left ventricle; MV, mitral valve. **B**, Representative echocardiogram views of the patients selected for iPSC generation and nuclear RNAseq profiling. FCA, four-chamber apical view; FCSC, four-chamber subcostal view; PLAX, parasternal long axis view. **C**, Average of LV lumen area/LV area ratio calculated from echocardiographic four-chamber apical views of the 3 patients selected for iPSC generation (HLHS iPSC) and the 4 patients selected for nuclear RNAseq profiling (HLHS RNAseq). Data are mean ± SEM.

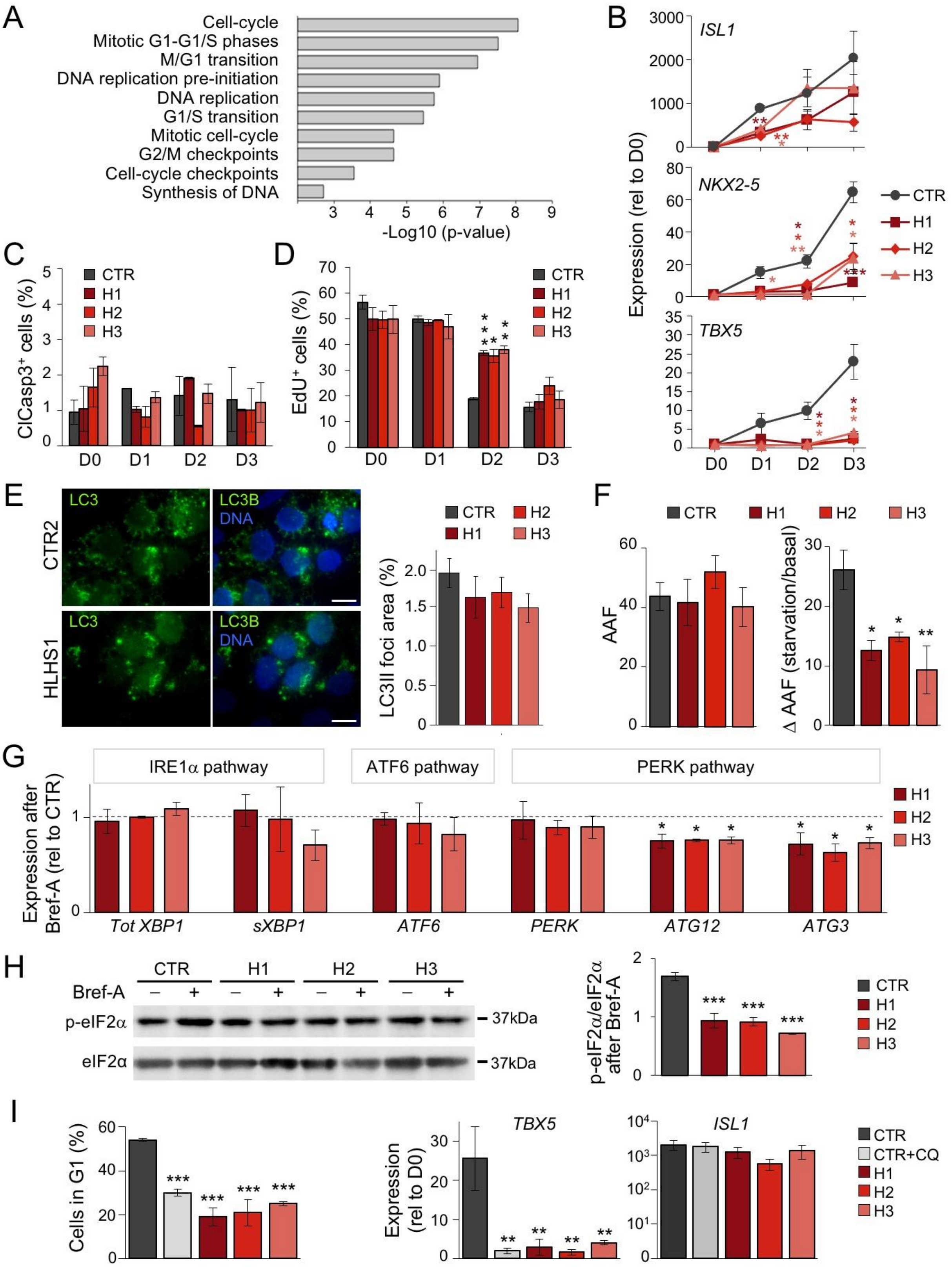


**Supplemental Figure VII. Cell-cycle, apoptosis, autophagy, and UPR analyses in HLHS and control CPs (Related to Figure 4). A**, Top-ten enriched GO pathways from cell-cycle DEGs of HLHS CPs at D3. **B**, Time-course qRT-PCR analysis of *ISL1*, *NKX2-5* and *TBX5*

during CP differentiation of HLHS and control iPSCs. Data are mean ± SEM, n=3-6 differentiations per line. *p<0.05, **p<0.01, ***p<0.001 compared to CTR (two-way ANOVA). **C**, Time-course immunostaining quantification of activated caspase-3 (ClCasp3) in HLHS and control cells. Data are mean ± SEM, n=3 differentiations per line, N=300 cells per line. **D**, Flow-cytometry quantification of $EdU^+$ cells in HLHS and control cells during CP differentiation. Data are mean ± SEM, n=3 differentiations per line, N≥20000 cells per sample at each time point. *p<0.05, **p<0.01 ***p<0.001 compared to CTR (two-way ANOVA). **E**, Immunofluorescence analysis of LC3II foci in HLHS and control CPs at D3. Scale bar, 5 μm. Data are mean ± SEM, N=30 cells from 3 independent differentiations per line. **F**, FACS-based quantification of the autophagy activity factor (AAF) in HLHS and control CPs at D3 by autophagic vesicle labeling with CYTO-ID. Data are mean ± SEM, n=3 differentiations per line, N≥20000 cells per sample. *p<0.05, **p<0.01 compared to CTR (one-way ANOVA). **G**, Expression levels of UPR actors in HLHS and control CPs at D3 after brefeldin-A treatment. Data are mean ± SEM, n=2-5 differentiations per line. *p<0.05 compared to CTR (one-way ANOVA). **H**, Western blot of total and phosphorylated eIF2a (eIF2a and p-eIF2a) in HLHS and control CPs at D3 with or without brefeldin-A. Data are mean ± SEM, n=9 (CTR), n=3 (HLHS2) and n=2 (HLHS1 and HLHS3) independent differentiations. ***p<0.001 compared to CTR (one-way ANOVA). **I**, Analysis of 12-hour chloroquine (CQ) treatment in control CPs. Left bar graph shows propidium iodide staining-based quantification of cells in G1 phase in HLHS and control CPs at D2 with or without chloroquine. Middle and right bar graphs summarize qRT-PCR analysis of *TBX5* and *ISL1* expression, respectively, in control CPs (with or without chloroquine) and in HLHS CPs at D3. Data are mean ± SEM, n=2-4 differentiations per line. N≥20000 cells per sample for G1 phase analysis. **p<0.01, ***p<0.001 compared to CTR (one-way ANOVA).

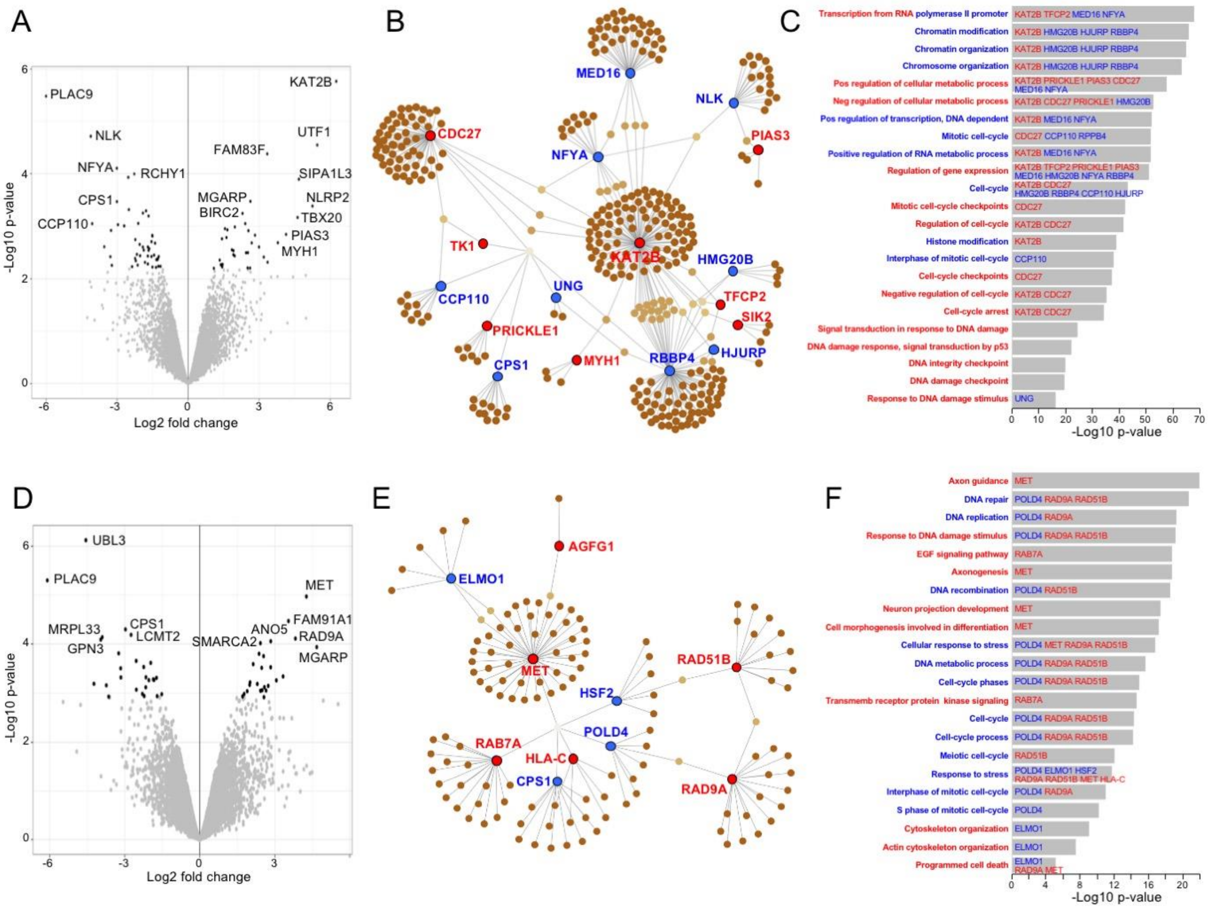


**Supplemental Figure VIII. Proteomic analysis of HLHS and control iPSC-CMs at D8 and D14 of differentiation. A** and **D**, Volcano plot of differentially expressed proteins (DEPs; 1.5-fold-expression, FDR <0.05) between HLHS and control CM at D8 (A) and D14 (D). **B** and **C**, Network analysis of DEPs at D8 (B) and the correspondent enriched GO terms (C). Upregulated and downregulated proteins and pathways are depicted in red and blue, respectively. **E** and **F**, Network analysis of DEPs at D14 (E) and the correspondent enriched GO terms (F). Upregulated and downregulated proteins and pathways are depicted in red and blue, respectively.

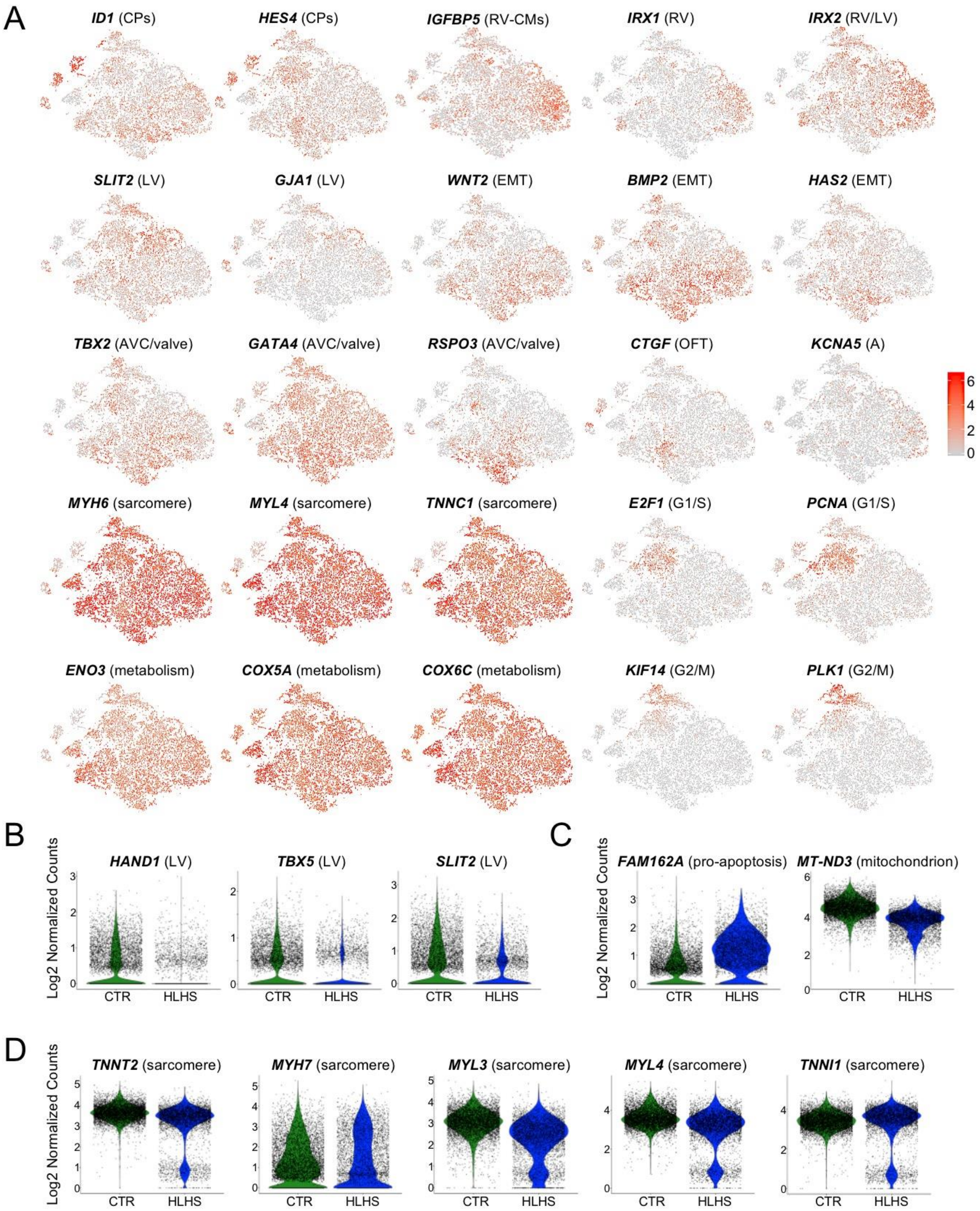


**Supplemental Figure IX. Single-cell RNAseq of HLHS and control iPSC-CMs at D14 of differentiation (Related to Figure 5). A**, Expression of selected genes marking subpopulations on t-SNE plot. EMT, epithelial-mesenchymal transition. **B**-**D**, Violin plots of selected DEGs between control and HLHS cells. All genes represented have a $p<0.05$.

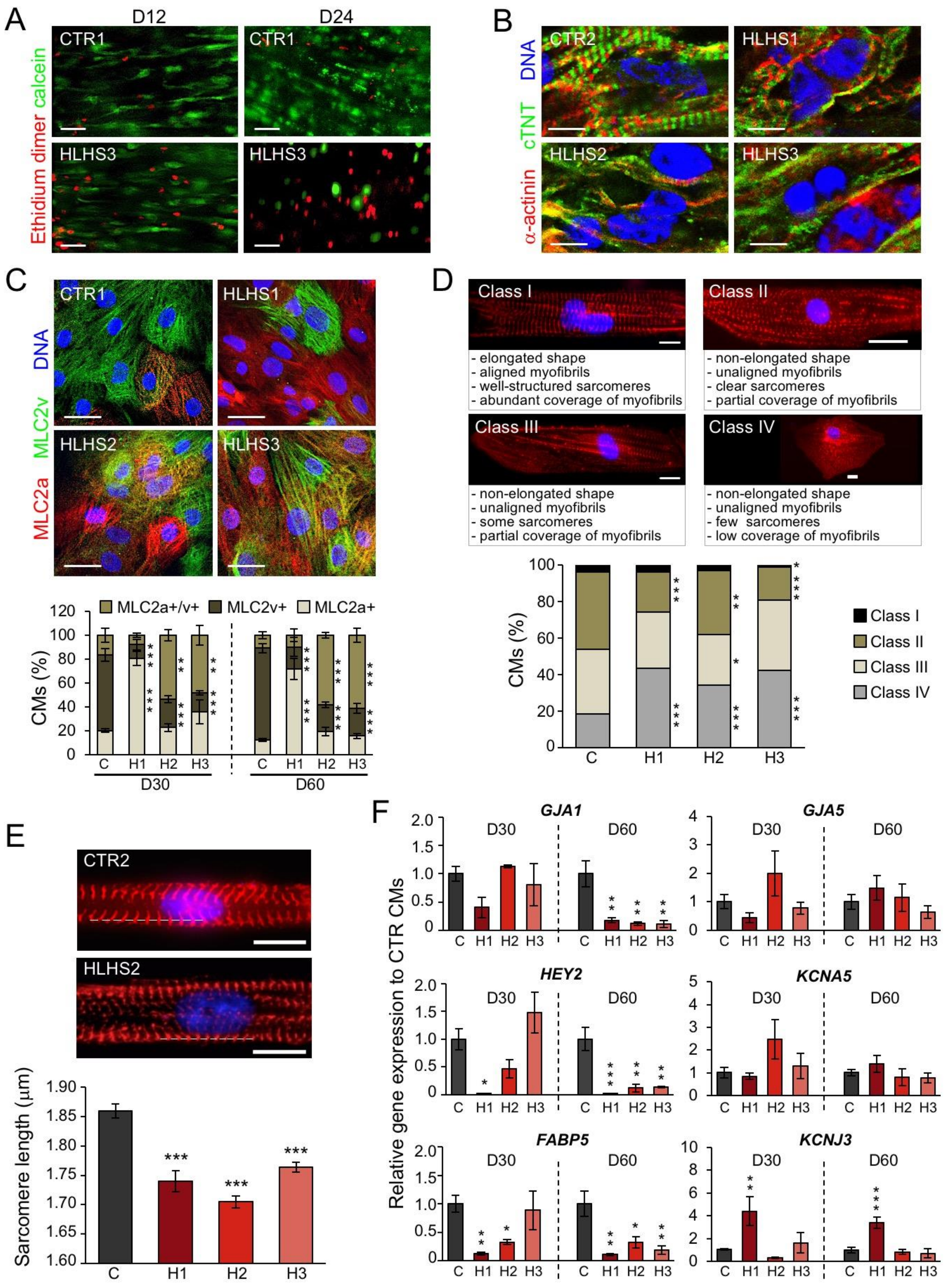


**Supplemental Figure X. HLHS CMs show increased cell death and sarcomeric disorganization in 3D patches as well as maturation defects in 2D culture (Related to Figure 7). A**, Representative staining with calcein and ethidium dimer of HLHS and control

3D heart patches at D12 and D24. Scale bar, 100 µm. **B**, Representative immunostaining for cTNT and α-actinin of HLHS and control 3D heart patches at D24. Scale bar, 10 µm. **C**, Top, representative immunostaining for MLC2a and MLC2v in HLHS and control CMs at D60 of 2D culture. Scale bar, 25 µm. Bottom, percentage of HLHS and control CMs expressing MLC2a, MLC2v or both markers at D30 and D60. Data are mean ± SEM, D30: N=1356 cells from 5 differentiations for CTR; N=764 cells for HLHS1, N=739 for HLHS2, N=876 for HLHS3 from 3 differentiations per line; D60: N=1475 cells from 6 differentiations for CTR; N=831 cells for HLHS1, N=781 for HLHS2, N=848 for HLHS3 from 3 differentiations per line. **p<0.01, ***p<0.001 compared to CTR (one-way ANOVA). **D**, Top, representative immunostaining for α-actinin labeling sarcomere Z-discs and description of sarcomere organization scoring in CMs at D60 of 2D culture on micropatterned slides to induce intracellular alignment of myofibrils. Class IV represents the most disarrayed sarcomeric organization. Scale bars, 20 µm. Bottom, percentage of CMs of individual sarcomeric structure classes in control and HLHS lines. n=348 CMs for CTR, n=355 for HLHS1, n=422 for HLHS2, n=335 for HLHS3 from 2 differentiations per line. *p<0.05, **p<0.01, ***p<0.001 compared to CTR (Chi-squared test). **E**, Analysis of sarcomere length in α-actinin-stained, class-I-scored control and HLHS CMs after treatment with the relaxant 2,3-butanedione 2-monoxime (BDM). Dotted line in images indicates sarcomere length calculated from intensity profile of line scans along myofibrils. Scale bars, 10 µm. Data are mean ± SEM, n=165 for CTR, n=60 for HLHS1, n=173 for HLHS2, and n=222 for HLHS3 cells from 2 differentiations per line. ***p<0.001 compared to CTR (Kruskal-Wallis test). **F**, Quantitative PCR gene expression analysis of ventricular (*GJA1, HEY2, FABP5*) and atrial (*GJA5, KCNA5, KCNJ3*) markers in HLHS and control CMs at D30 and D60. Data are mean ± SEM from 3-7 differentiations per line. *p<0.05, **p<0.01, ***p<0.001 compared to CTR (one-way ANOVA).

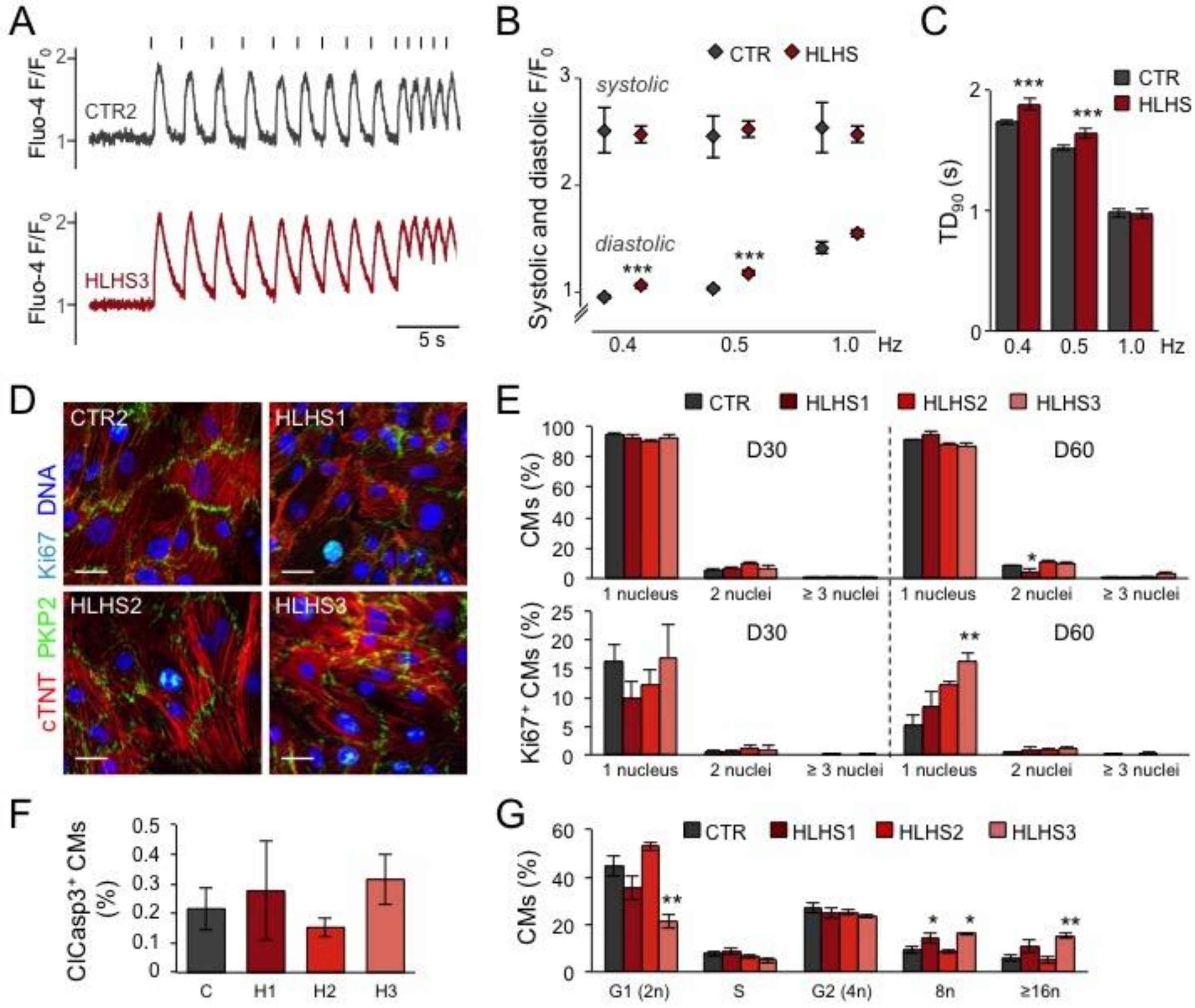


**Supplemental Figure XI. HLHS CMs in 2D culture show defective intracellular $Ca^{2+}$ cycling but no aberrant multinucleation or apoptosis (Related to Figure 7). A**, Representative images of Fluo-4-based intracellular calcium transients from single control and HLHS D30 CMs in 2D at increasing pacing rates (0.4, 0.5, and 1.0 Hz). Fluo-4 fluorescence was normalized to the initial fluorescence level (F/F0). Vertical bars over the tracings represent the stimulation pulses. **B** and **C**, Mean systolic and diastolic $Ca^{2+}$ levels (expressed as F/F0, B) and mean $Ca^{2+}$ transient duration at 90% decay ($TD_{90}$, C) in control and HLHS D30 CMs at the indicated pacing rates. Data are mean ± SEM, n=29 (CTR) and 90 (HLHS) CMs from 3 independent differentiations. ***p<0.001 compared to CTR at the same pacing rate (Mann Whitney test). **D**, Representative immunostaining for cTNT, PKP2, Ki67 in HLHS and control CMs at D60 in 2D culture. Scale bars, 25 µm. **E**, Percentage of

HLHS and control CMs with 1, 2 and ≥3 nuclei at D30 and D60 (top). Percentage of $Ki67^+$ HLHS and control CMs with 1, 2 and ≥3 nuclei at D30 and D60 (bottom). Data are mean ± SEM, D30: N=1414 CMs for CTR from 5 differentiations; N=978 CMs for HLHS1, N=680 for HLHS2 from 3 differentiations per line; N=980 for HLHS3 from 4 differentiations; D60: N=1490 CMs for CTR from 4 differentiations; N=1306 CMs for HLHS1 and N=901 for HLHS2 from 3 differentiations per line, N=1229 for HLHS3 from 4 differentiations. *p<0.05, **p<0.01 compared to CTR (one-way ANOVA). **F**, Percentage of $ClCasp3^+$ HLHS and control CMs at D60, assessed by flow cytometry. Data are ± SEM; n=7 (CTR), n=3 (HLHS1), and n=4 (HLHS2 and HLHS3) independent differentiations; N≥20000 cells per sample. **G**, Quantification of DNA content by propidium iodide flow cytometry analysis of HLHS and control CMs at D60. Data are ± SEM; n=5 (CTR), n=4 (HLHS1), n=5 (HLHS2), and n=3 (HLHS3) differentiations; N≥20000 cells per sample. *p<0.05, **p<0.01 compared to CTR (one-way ANOVA).

#### LEGENDS OF SUPPLEMENTAL EXCEL TABLES

**Supplemental Table I. Clinical and demographic information of individuals selected for WES, WES results, and data source for WES analyses (Related to Figure 1 and Supplemental Figure II).**

**Supplemental Table II. Clinical and demographic information of individuals selected for nuclear RNAseq (nRNAseq) of native ventricle CMs, nRNAseq results, and data source for nRNAseq analyses (Related to Figure 2 and Supplemental Figure III).**

**Supplemental Table III. Clinical and demographic information of individuals selected for iPSC line generation and data source for RNAseq, single-cell RNAseq, and proteomic analyses on iPSC-derived cardiac cells (Related to Figures 3 and 5 and Supplemental Figures VII and VIII).**

**Supplemental Table IV. Data source for quantifications and statistics**
